# Automated construction of cognitive maps with visual predictive coding

**DOI:** 10.1101/2023.09.18.558369

**Authors:** James A. Gornet, Matt Thomson

## Abstract

Humans construct internal cognitive maps of their environment directly from sensory inputs without access to a system of explicit coordinates or distance measurements. While machine learning algorithms like SLAM utilize specialized inference procedures to identify visual features and construct spatial maps from visual and odometry data, the general nature of cognitive maps in the brain suggests a unified mapping algorithmic strategy that can generalize to auditory, tactile, and linguistic inputs. Here, we demonstrate that predictive coding provides a natural and versatile neural network algorithm for constructing spatial maps using sensory data. We introduce a framework in which an agent navigates a virtual environment while engaging in visual predictive coding using a self-attention-equipped convolutional neural network. While learning a next image prediction task, the agent automatically constructs an internal representation of the environment that quantitatively reflects spatial distances. The internal map enables the agent to pinpoint its location relative to landmarks using only visual information.The predictive coding network generates a vectorized encoding of the environment that supports vector navigation where individual latent space units delineate localized, overlapping neighborhoods in the environment. Broadly, our work introduces predictive coding as a unified algorithmic framework for constructing cognitive maps that can naturally extend to the mapping of auditory, sensorimotor, and linguistic inputs.

---

Space and time are fundamental physical structures in the natural world, and all organisms have evolved strategies for navigating space to forage, mate, and escape predation.^1–3^. In humans and other mammals, the concept of a spatial or cognitive map has been postulated to underlie spatial reasoning tasks^4–6^. A spatial map is an internal, neural representation of an animal’s environment that marks the location of landmarks, food, water, shelter, and then can be queried for navigation and planning. The neural algorithms underlying spatial mapping are thought to generalize to other sensory modes to provide cognitive representations of auditory and somatosensory data^7^ as well as to construct internal maps of more abstract information including concepts^8,9^, tasks^10^, semantic information^11–13^, and memories^14^. Empirical evidence suggest that the brain uses common cognitive mapping strategies for spatial and non-spatial sensory information so that common mapping algorithms might exist that can map and navigate over not only visual but also semantic information and logical rules inferred from experience^7,8,15^. In such a paradigm reasoning itself could be implemented as a form of navigation within a cognitive map of concepts, facts, and ideas.

Since the notion of a spatial or cognitive map emerged, the question of how environments are represented within the brain and how the maps can be learned from experience has been a central question in neuroscience^16^. Place cells in the hippocampus are neurons that are active when an animal transits through a specific location in an environment^16^. Grid cells in the entorhinal cortex fire in regular spatial intervals and likely track an organism’s displacement in the environment^17,18^. Yet with the identification of a substrate for the representation of space, the question of how a spatial map can be learned from sensory data has remained, and the neural algorithms that enable the construction of spatial and other cognitive maps remain poorly understood.

Empirical work in machine learning has demonstrated that deep neural networks can solve spatial navigation tasks as well as perform path prediction and grid cell formation^19,20^. Cueva & Wei^19^ and Banino *et al*.^20^ demonstrate that neural networks can learn to perform path prediction and that networks generate firing patterns that resemble the firing patterns of grid cells in the entorhinal cortex. Crane *et al*.^21^, Zhang *et al*.^22^, and Banino *et al*.^20^ demonstrate navigation algorithms that require the environment’s map or using firing patterns that resemble place cells in the hippocampus. These studies allow an agent to access environmental coordinates explicitly^19^ or initialize a model with place cells that represent specific locations in an arena^20^. In machine learning and autonomous navigation, a variety of algorithms have been developed to perform mapping tasks including SLAM and monocular SLAM algorithms^23–26^ as well as neural network implementations^27–29^. Yet, SLAM algorithms contain many specific inference strategies, like visual feature and object detection, that are specifically engineered for map building, wayfinding, and pose estimation based on visual information. Whereas extensive research in computer vision and machine learning use video frames, these studies do not extract representations of the environment’s map^30,31^. A unified theoretical and mathematical framework for understanding the mapping of spaces based on sensory information remains incomplete.

Predictive coding has been proposed as a unifying theory of neural function where the fundamental goal of a neural system is to predict future observations given past data^32–34^. When an agent explores a physical environment, temporal correlations in sensory observations reflect the structure of the physical environment. Landmarks nearby one another in space will also be observed in temporal sequence. In this way, predicting observations in a temporal series of sensory observations requires an agent to internalize some implicit information about a spatial domain. Historically, Poincare motivated the possibility of spatial mapping through a predictive coding strategy where an agent assembles a global representation of an environment by gluing together information gathered through local exploration^35,36^. The exploratory paths together contain information that could, in principle, enable the assembly of a spatial map for both flat and curved manifolds. Indeed, extended Kalman filters^25,37^ for SLAM perform a form of predictive coding by directly mapping visual changes and movement to spatial changes. However, extended Kalman filters as well as other SLAM approachs require intricate strategies for landmark size calibration, image feature extraction, and models of the camera’s distortion whereas biological systems can solve flexible mapping and navigation that engineered systems cannot. Yet, while the concept of predictive coding for spatial mapping is intuitively attractive, a major challenge is the development of algorithms that can glue together local, sensory information gathered by an agent into a global, internally consistent environmental map. Connections between mapping and predictive coding in the literature have primarily focused on situations where an agent has explicit access to its spatial location as a state variable^38–40^. The problem of building spatial maps *de novo* from sensory data remains poorly understood.

Here, we demonstrate that a neural network trained on a sensory, predictive coding task can construct an implicit spatial map of an environment by assembling observations acquired along local exploratory paths into a global representation of a physical space within the network’s latent space. We analyze sensory predictive coding theoretically and demonstrate mathematically that solutions to the predictive sensory inference problem have a mathematical structure that can naturally be implemented by a neural network with a ‘path-encoder,’ an internal spatial map, and a ‘sensory decoder,’ and trained using backpropagation. In such a paradigm, a network learns an internal map of its environment by inferring an internal geometric representation that supports predictive sensory inference. We implement sensory predictive coding within an agent that explores a virtual environment while performing visual predictive coding using a convolutional neural network with self-attention. Following network training during exploration, we find that the encoder network embeds images collected by an agent exploring an environment into an internal representation of space. Within the embedding, the distances between images reflect their relative spatial position, not object-level similarity between images. During exploratory training, the network implicitly assembles information from local paths into a global representation of space as it performs a next image inference problem. Fundamentally, we connect predictive coding and mapping tasks, demonstrating a computational and mathematical strategy for integrating information from local measurements into a global self-consistent environmental model.

## Mathematical formulation of spatial mapping as sensory predictive coding

In this paper, we aim to understand how a spatial map can be assembled by an agent that is making sensory observations while exploring an environment. Papers in the literature that study connections between predictive coding and mapping have primarily focused on situations where an agent has access to its ‘state’ or location in the environment^38–40^. Here, we develop a theoretical model and neural network implementation^†^ of sensory predictive coding that illustrates why and how an internal spatial map can emerge naturally as a solution to sensory inference problems. We, first, formulate a theoretical model of visual predictive coding and demonstrate that the predictive coding problem can be solved by an inference procedure that constructs an implicit representation of an agent’s environment to predict future sensory observations. The theoretical analysis also suggests that the underlying inference problem that can be solved by an encoder-decoder neural network that infers spatial position based upon observed image sequences.

We consider an agent exploring an environment, Ω ⊂ ℝ^2^, while acquiring visual information in the form of pixel valued image vectors *I*_*x*_ ∈ ℝ^*m*×*n*^ given an *x*∈Ω. The agent’s environment Ω is a bounded subset of ℝ^2^ that could contain obstructions and holes. In general, at any given time, *t*, the agent’s state can be characterized by a position *x*(*t*) and orientation *θ* (*t*) where *x* (*t*) and *θ* (*t*) are coordinates within a global coordinate system unknown to the agent.

The agent’s environment comes equipped with a visual scene, and the agent makes observations by acquiring image vectors 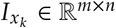 as it moves along a sequence of points *x*_*k*_. At every position *x* and orientation *θ*, the agent acquires an image by effectively sampling from an image the conditional probability distribution *P* (*I*|*x*_*k*_, *θ*_*k*_) which encodes the probability of observing a specific image vector *I* when the agent is positioned at position *x*_*k*_ and orientation *θ*_*k*_. The distribution *P* (*I*|*x*), *θ* has a deterministic and stochastic component where the deterministic component is set by landmarks in the environment while stochastic effects can emerge due to changes in lighting, background, and scene dynamics. Mathematically, we can view *P* (*I*|*x, θ*) as a function on a vector bundle with base space Ω and total space Ω × *I*^43^. The function assigns an observation probability to every possible image vector for an agent positioned at a point (*x, θ*). Intuitively, the agent’s observations preserve the geometric structure of the environment: the spatial structure influences temporal correlations.

In the predictive coding problem, the agent moves along a series of points (*x*_0_, *θ*_0_), (*x*_1_, *θ*_1_), …, (*x*_*k*_, *θ*_*k*_) while acquiring images *I*_0_, *I*_1_, … *I*_*k*_. The motion of the agent in Ω is generated by a Markov process with transition probabilities *P*(*x*_*i*+1_, *θ*_*i*+1_| *x*_*i*_, *θ*_*i*_). Note that the agent has access to the image observations *I*_*i*_ but not the spatial coordinates (*x*_*i*_, *θ*_*i*_). Given the set {*I*_0_ … *I*_*k*_ } the agent aims to predict *I*_*k*+1_. Mathematically, the image prediction problem can be solved theoretically through statistical inference by (a) inferring the posterIor probability distribution *P*(*I*_*k*+1_ |*I*_0_, *I*_1_….*I*_*k*_) from observations. Then, (b) given a specific sequence of observed images {*I*_0_ … *I*_*k*_ }, the agent can predict the next image *I*_*k* + 1_ by finding the image *I*_*k*+1_ that maximizes the posterior probability distribution *P*(*I*_*k*+1_|*I*_0_, *I*_1_ *I*_*k*_).

The posterior probability distribution *P*(*I*_*k*+1_ |*I*_0_, *I*_1_, ….*I*_*k*_) is by definition

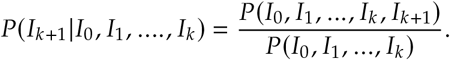

If we consider *P* (*I*_0_, *I*_1_ … *I*_*k*_, *I*_*k*+1_)to be a function of an implicit set of spatial coordinates (*x*_*i*_, *θ*_*i*_) where the (*x*_*i*_, *θ*_*i*_) provide an internal representation of the spatial environment. Then, we can express the posterior probability *P* (*I*_*k+1*_ *I*_0_, *I*_1_….*I*_*k*_) in terms of the implicit spatial representation

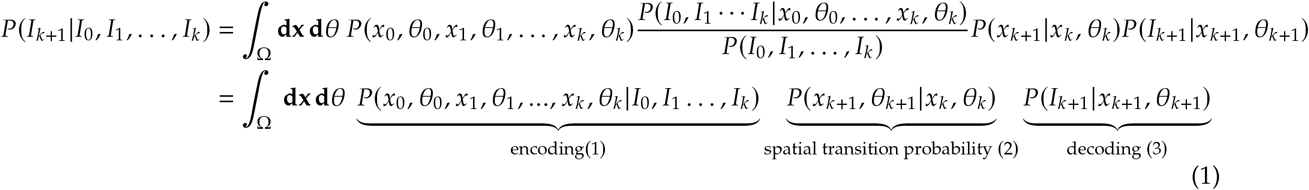

where in Equation 1 the integration is over all possible paths {(*x*_0_, *θ*_0_)…(*x*_*k*_, *θ*_*k*_)} in the domain Ω, **dx** = *dx*_0_ … *dx*_*k*_, and **d***θ* = *dθ*_0_ … *dθ*_*k*_. Equation 1 can be interpreted as a path integral over the domain Ω. The path integral assigns a probability to every possible path in the domain and then computes the probability that the agent will observe a next image *I*_*k*_ given an inferred location (*x*_*k*+ 1_, *θ*_*k*+1_). In detail, term (1) assigns a probability to every discrete path {(*x*_0_, *θ*_0_ … *x*_*k*_, *θ*_*k*_)}∈ Ω as the conditional likelihood of the path given the observed sequences of images {*I*_0_ … *I*_*k*_}. Term (2) computes the probability that an agent at a terminal position *x*_*k*_ moves to the position (*x*_*k+1*_, *θ*_*k*+ 1_) given the Markov transition function *P* (*x*_*k*+1_, *θ*_*k*+1_ |*x*_*k*_, *θ*_*k*_). Term (3) is the conditional probability that image *I*_*k*+1_ is observed given that the agent is at position (*x*_*k*+1_, *θ*_*k*+1_).

Conceptually, the product of terms solves the next image prediction problem in three steps. First (1), estimating the probability that an agent has traversed a particular sequence of points given the observed images; second (2), estimating the next position of the agent (*x*_*k*+1_, *θ*_*k*+1_) for each potential path; and third (3), computing the probability of observing a next image *I*_*k*_+_1_ given the inferred terminal location *x*_*k* + 1_ of the agent. Critically, an algorithm that implements the inference procedure encoded in the equation would construct an internal but implicit representation of the environment as a coordinate system **x, *θ*** that is learned by the agent and used during the next image inference procedure. The coordinate system provides an internal, inferred representation of the agent’s environment that is used to estimate future image observation probabilities. Thus, our theoretical framework demonstrates how an agent might construct an implicit representation of its spatial environment by solving the predictive coding problem.

The three step inference procedure represented in the equation for *P*(*I*_*k*+1_ |*I*_0_ … *I*_*k*_) can be directly implemented in a neural network architecture, as demonstrated in the Appendix. The first term acts as an ‘encoder’ network that computes the probability that the agent has traversed a path {(*x*_0_, *θ*_0_)…( *x*_*k*_, *θ*_*k*_)} given an observed image sequence *I*_0_, …, *I*_*k*_ that has been observed by the network (Figure 1(**b**)). The network can, then, estimate the next position of the agent (*x*_*k*_+_1_, *θ*_*k*_+_1_) given an inferred location (*x*_*k*_, *θ*_*k*_), and apply a decoding network to compute *P* (*I*_*k*+1_ |*x*_*k*+1_, *θ*_*k+1*_ while outputting the prediction *I*_*k*+1_ using a decoder. A network trained through visual experience must learn an internal coordinate system and representation **x, *θ*** that not only offers an environmental representation but also establishes a connection between observed images *I*_*j*_ and inferred locations (*x*_*j*_, *θ*_*j*_).

**Figure 1.**
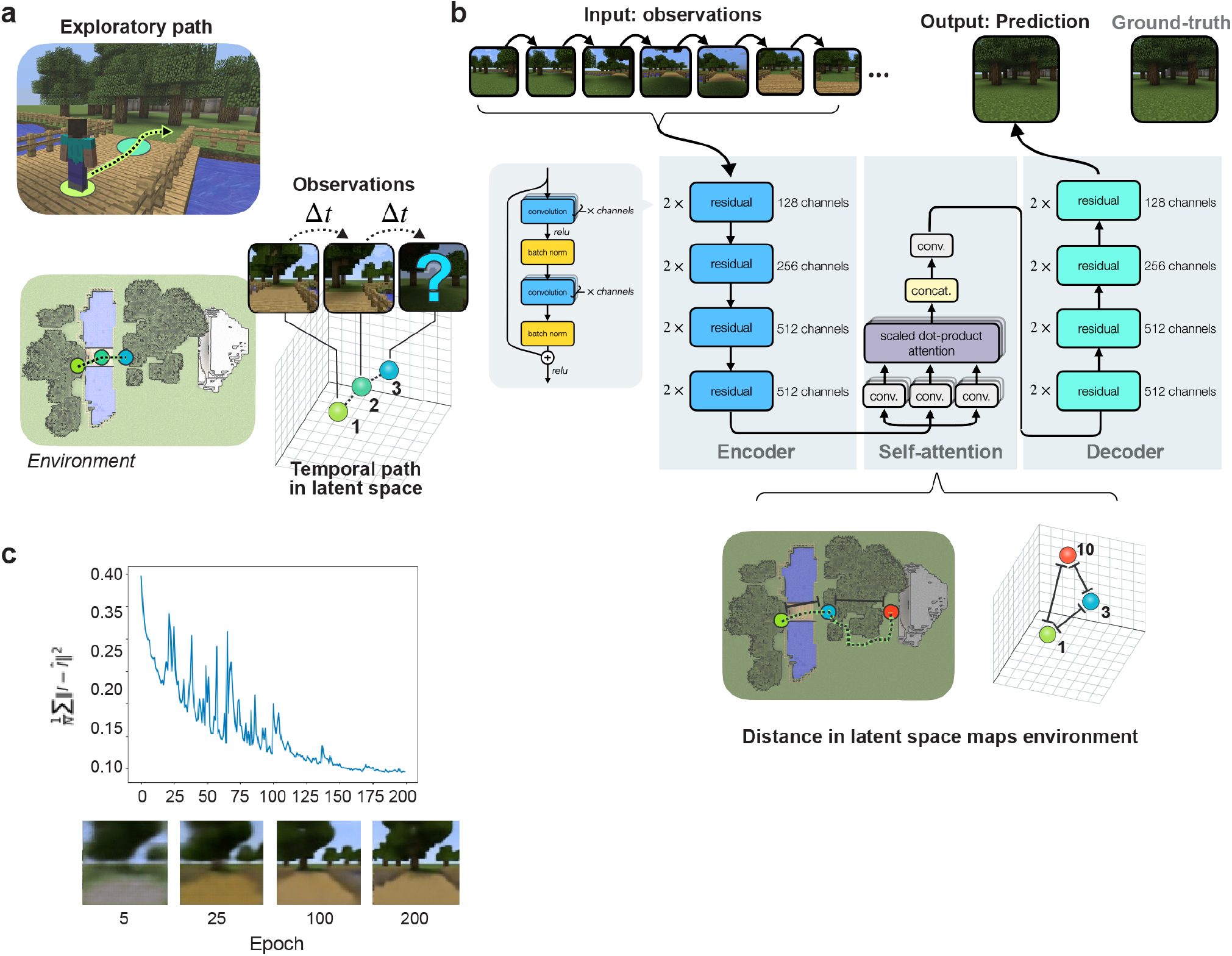
A predictive coding neural network explores a virtual environment. In predictive coding, a model predicts observations and updates its parameters using the prediction error. **a**, an agent’s traverses its environment by taking the most direct path to random positions. **b**, a self-attention-based encoder-decoder neural network architecture learns to perform predictive coding. A ResNet-18 convolutional neural network acts as an encoder; self-attention is performed with 8 heads, and a corresponding ResNet-18 convolutional neural network performing decoding to the predicted image.**c**, the neural network learns to perform predictive coding effectively—with a mean-squared error of 0.094 between the actual and predicted images.

### A neural network performs accurate predictive coding within a virtual environment

Motivated by the implicit representation of space contained in the predictive coding inference problem, we developed a computational implementation of a predictive coding agent, and studied the representation of space learned by that agent as it explored a virtual environment. We first create an environment with the Malmo environment in Minecraft^44^. The physical environment measures 40 × 65 lattice units and encapsulates three aspects of visual scenes: a cave provides a global visual landmark, a forest provides degeneracy between visual scenes, and a river with a bridge constrains how an agent traverses the environment (Figure 1(**a**)). An agent follows paths (Figure S5(**b-c**)), determined by *A*^*^ search, between randomly sampled positions and receives visual images along every path.

To perform predictive coding, we construct an encoder-decoder convolutional neural network (CNN) with a ResNet-18 architecture^45^ for the encoder and a corresponding ResNet-18 architecture with transposed convolutions in the decoder (Figure 1(**b**)). The encoder-decoder architecture uses the U-Net architecture^46^ to pass the encoded latent units into the decoder. Multi-headed attention^47^ processes the sequence of encoded latent units to encode the history of past visual observations. The multi-headed attention has *h* = 8 heads. For the encoded latent units with dimension *D* = *C*×*H*×*W*, the dimension *d* of a single head is *d* = *C* × *H* × *W*/*h*.

The predictive coder approximates predictive coding by minimizing the mean-squared error between the actual observation and its predicted observation. The predictive coder trains on 82, 630 samples for 200 epochs with gradient descent optimization with Nesterov momentum^48^, a weight decay of 5 ×10^−6^, and a learning rate of 10^−1^ adjusted by OneCycle learning rate scheduling^49^. The optimized predictive coder has a mean-squared error between the predicted and actual images of 0.094 and a good visual fidelity (Figure 1(**c**)).

### Predictive coding network constructs an implicit spatial map

We show that the predictive coder creates an implicit spatial map by demonstrating it recovers the environment’s spatial position and distance. We encode the image sequences using the predictive coder’s encoder to analyze the encoded sequence as the predictive coder’s latent units. To measure the positional information in the predictive coder, we train a neural network to predict the agent’s position from the predictive coder’s latent units (Figure 1(**a**)). The neural network’s prediction error

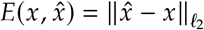

indirectly measures the predictive coder’s positional information. To provide comparative baselines, we construct a position prediction model. To lower bound the prediction error, we construct a model that gives the agent’s actual position with small additive Gaussian noise

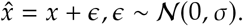

To compare the predictive coder to the baselines, we compare the prediction error histograms (Figure 2(**b**)).

**Figure 2.**
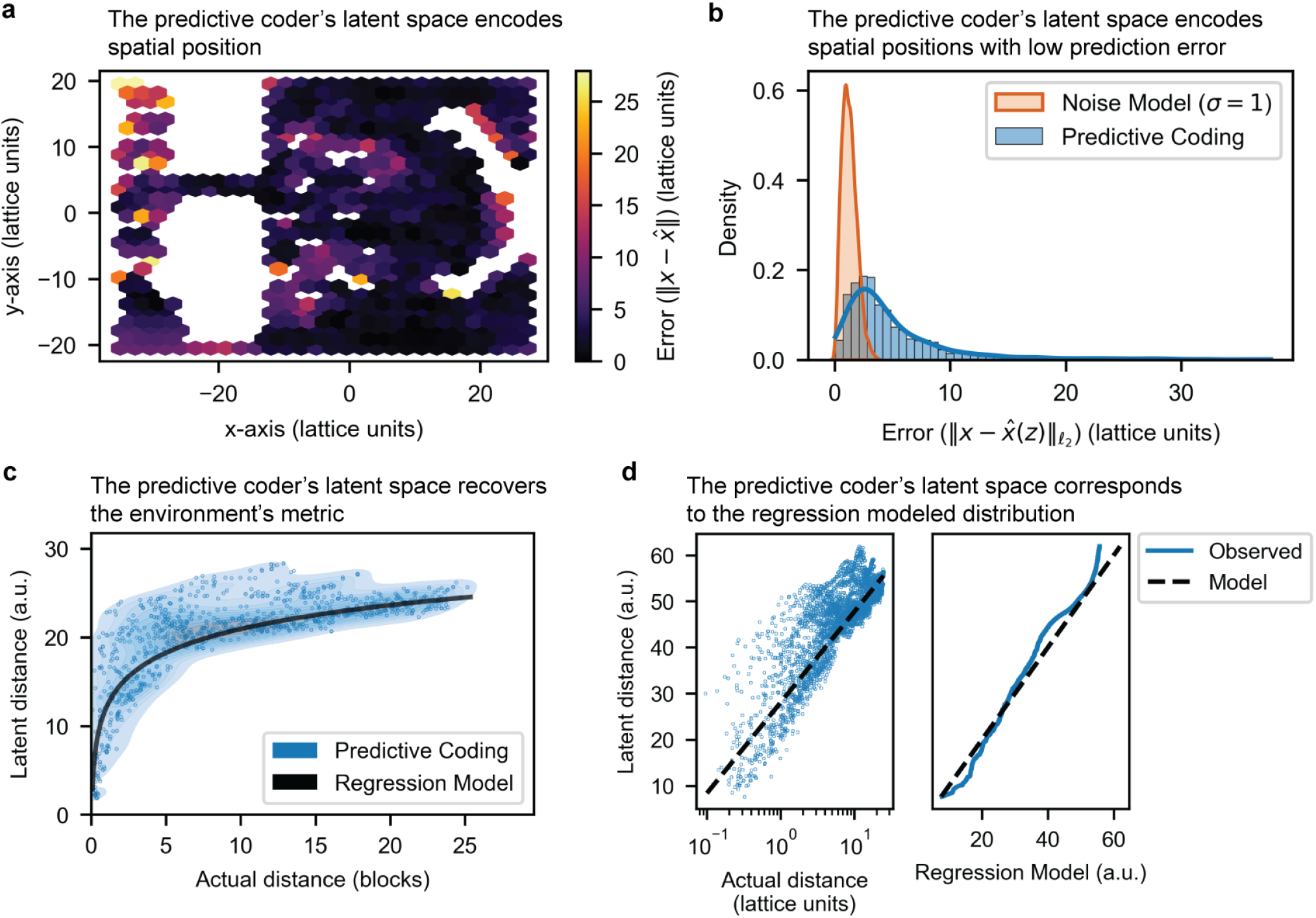
Predictive coding neural network constructs an implicit spatial map. **a-b**, The predictive coder’s latent space encodes accurate spatial positions. A neural network predicts the spatial location from the predictive coding’s latent space. **a**, a heatmap of the prediction errors between the actual position and the predictive coder’s predicted positions show a low prediction error. **b**, The histogram of prediction errors of positions from the predictive coder’s latent space show a low prediction error. As a baseline (Noise model (σ = 1 lattice unit)), actual positions with a small noise displacement gives an error model. **c**, predictive coding’s latent distances recover the environment’s spatial metric. Sequential visual images are mapped to the neural network’s latent space, and the latent space distances (*ℓ*_2_) are plotted with physical distances onto a joint density plot. An nonlinear regression model

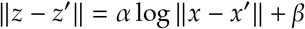

is shown as a baseline. **d**, a correlation plot and a quantile-quantile plot show the overlap between the empirical and model distributions.

The predictive coder encodes the environment’s spatial position to a low prediction error (Figure 2.d). The predictive coder has a mean error of 5.04 lattice units and > 80% of samples have an error < 7.3 lattice units. The additive Gaussian model with σ = 4 has a mean error of 4.98 lattice units and > 80% of samples with an error < 7.12 lattice units.

We show the predictive coder’s latent space recovers the local distances between the environment’s physical positions. For every path that the agent traverses, we calculate the local pairwise distances in physical space and in the predictive coder’s latent space with a neighborhood of 100 time points. To determine whether latent space distances correspond to physical distances, we calculate the joint density between latent space distances and physical distances (Figure 2(**c**)). We model the latent distances by fitting the physical distances with additive Gaussian noise to a logarithmic function

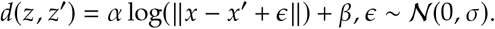

The modeled distribution is concentrated with the predictive coder’s distribution (Figure 2(**d**)) with a Pearson correlation coefficient of 0.827 and a Kullback-Leibler divergence (ⅅ_KL_(*p*_PC_∥*p*_model_)) of 0.429 bits.

### Predictive coding network learns spatial proximity not image similarity

In the previous section, we show that a neural network that performs predictive coding learns an internal representation of its physical environment within its latent space. Here, we demonstrate that the prediction task itself is essential for spatial mapping. Prediction forces a network to learn spatial proximity and not merely image similarity. Many frameworks including principal components analysis, IsoMap^50^, and autoencoder neural networks can collocate images by visual similarity. While similar scenes might be proximate in space, similar scenes can also be spatially divergent. For example, the virtual environment we constructed has two different ‘forest’ regions that are separated by a lake. Thus, in the two forest environments might generate similar images but are actually each closer to the lake region than to one another (Figure 1(**a**)).

To demonstrate the central role for prediction in mapping, we compared the latent representation of images generated by the predictive coding network to a representation learned by an autoencoder. The auto-encoder network has a similar architecture to the predictive encoder but encodes a *single* image observation in a latent space, and decodes the same observations. As the auto-encoder only operates on a single image— rather than a sequence, the auto-encoder learns an embedding based on image proximity not underlying spatial relationships. As with the predictive coder, the auto-encoder (Figure 3(**a**)) trains to minimize the mean-squared error between the actual image and the predicted image on 82, 630 samples for 200 epochs with gradient descent optimization with Nesterov momentum, a weight decay of 5×10^−6^, and a learning rate of 10^−1^ adjusted by the OneCycle learning rate scheduler. The auto-encoder has mean-squared error of 0.039 and a high visual fidelity.

**Figure 3.**
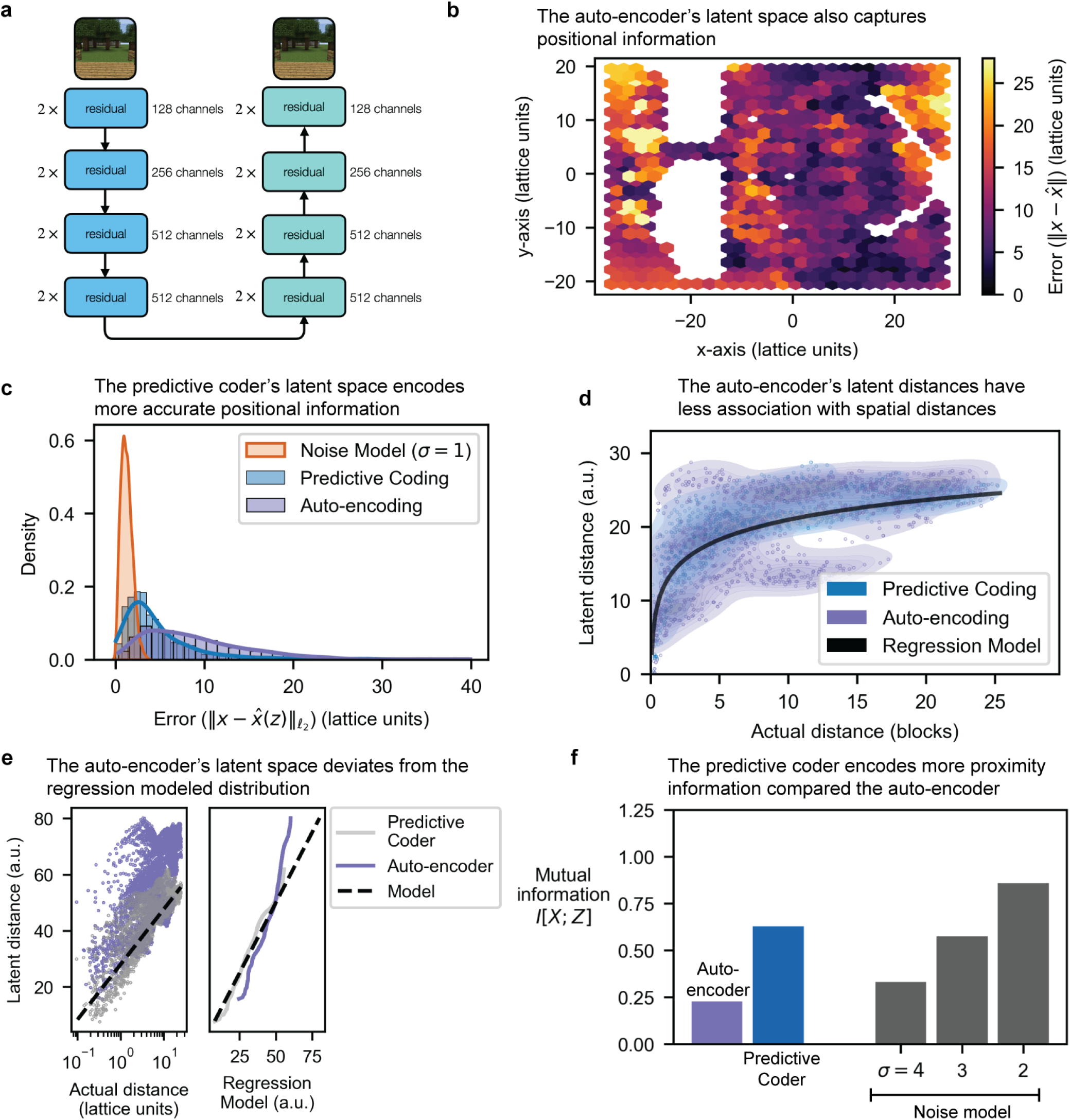
Predictive coding network learns spatial proximity not image similarity. **a**, an autoencoding neural network compresses visual images into a low-dimensional latent vector and reconstructs the image from the latent space. Auto-encoder trains on visual images from the environment *without any sequential order*. **b-c**, auto-encoding encodes lower resolution in positional information. **b**, a neural network predicts the spatial location from the auto-encoding’s latent space. A heatmap of the prediction errors between the actual position and the auto-encoder’s predicted positions show a higher prediction error—compared to the predictive coder. **c**, auto-encoding captures less positional information compared to predictive coding. The histogram shows the prediction errors of positions from the latent space of both the auto-encoder and the predictive coder. **d**, latent distances, however, show a weaker relationship with physical distances, as the joint histogram between physical and latent distances is less concentrated. **e**, a correlation plot and a quantile-quantile plot show a lower correlation and a lower density overlap between the empirical and model distributions. **f**, predictive coding’s latent units communicate more fine-grained spatial distances whereas auto-encoding communicates broad spatial regions. Joint density plots show the association between latent distances and physical distances for both predictive coding and auto-encoding. Predictive coding’s latent distances increase with spatial distances, with a higher concentration compared to auto-encoding.

The predictive coder encodes a higher resolution and more accurate spatial map in its latent space than the auto-encoder. As with the predictive coder, we train an auxiliary neural network to predict the agent’s position from the auto-encoder’s latent units (Figure 3(**b**)). The neural network’s prediction error indirectly measures the auto-encoder positional information. For greater than 80% of the auto-encoder’s points, its prediction error is less than 13.1 lattice units, as compared to the predictive coder that has > 80% of its samples below a prediction error of 7.3 lattice units (Figure 3(**c**)).

We also show that the predictive coder recovers the environment’s spatial distances with finer resolution compared to the auto-encoder. As with the predictive coder, we calculate the local pairwise distances in physical space and in the auto-encoder’s latent space, and we generate the joint density between the physical and latent distances (Figure 3(**d**)). Compared to the predictive coder’s joint density, the auto-encoder’s latent distances increase with the agent’s physical distance. The auto-encoder’s joint density shows a larger dispersion compared to the predictive coder’s joint density, indicating that the auto-encoder encodes spatial distances with higher uncertainty.

We can quantitatively measure the dispersion in the auto-encoder’s joint density by calculating mutual information of the joint density (Figure 3(**e**))

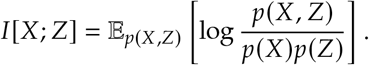

The auto-encoder has a mutual information of 0.227 bits while the predictive coder has a mutual information of 0.627 bits. As a comparison, positions with additive Gaussian noise having a standard deviation σ of 2 lattice units has a mutual information of 0.911 bits. The predictive coder encodes 0.400 additional bits of distance information to the auto-encoder. The predictive coder’s additional distance information of 0.4 bits exceeds the auto-encoder’s distance information of 0.227 bits, which indicates the temporal dependencies encoded by the predictive coder capture more spatial information compared to visual similarity.

### Predictive coding network maps visual degenerate environments whereas auto-encoding cannot

The sequential prediction task is beneficial for spatial mapping: the predictive coder captures more accurate spatial information compared to the auto-encoder, and predictive coder’s latent distances have a stronger correspondence to the environment’s metric. However, it is unclear whether predictive coding is *necessary* (as opposed to *beneficial*) to recover an environment’s map; an auto-encoder may still recover the environment’s map. In this section, we demonstrate that predictive coding is necessary for recovering an environment’s map. First, we show empirically that there exist environments that auto-encoding cannot recover. Second, we provide insight into why the auto-encoder fails with a theorem showing that auto-encoding cannot recover many environments—specifically, environments with visually similar yet spatially different locations.

In the previous sections, the agent explores a natural environment with forest, river, and cave landmarks. While this environment models exploration in outdoor environments, the lack of controlled visual scenes complicates interpreting predictive coder and auto-encoder. We introduce a circular corridor (Figure 4(**a**)) to introduce visual scenes that *visually identical*—rather than *visually similar*—yet spatially different. Specifically, the rooms revolve clockwise as red, green, red, blue, and yellow; there exist two distinct red rooms. The two distinct red rooms answers two questions: can the the predictive coder and auto-encoder recover the map for environments with visual symmetry, and does the predictive coder recover a global map or a relative map? In other words, does the predictive coder recovers the circular corridor’s geometry, or does it learn a linear hallway?

**Figure 4.**
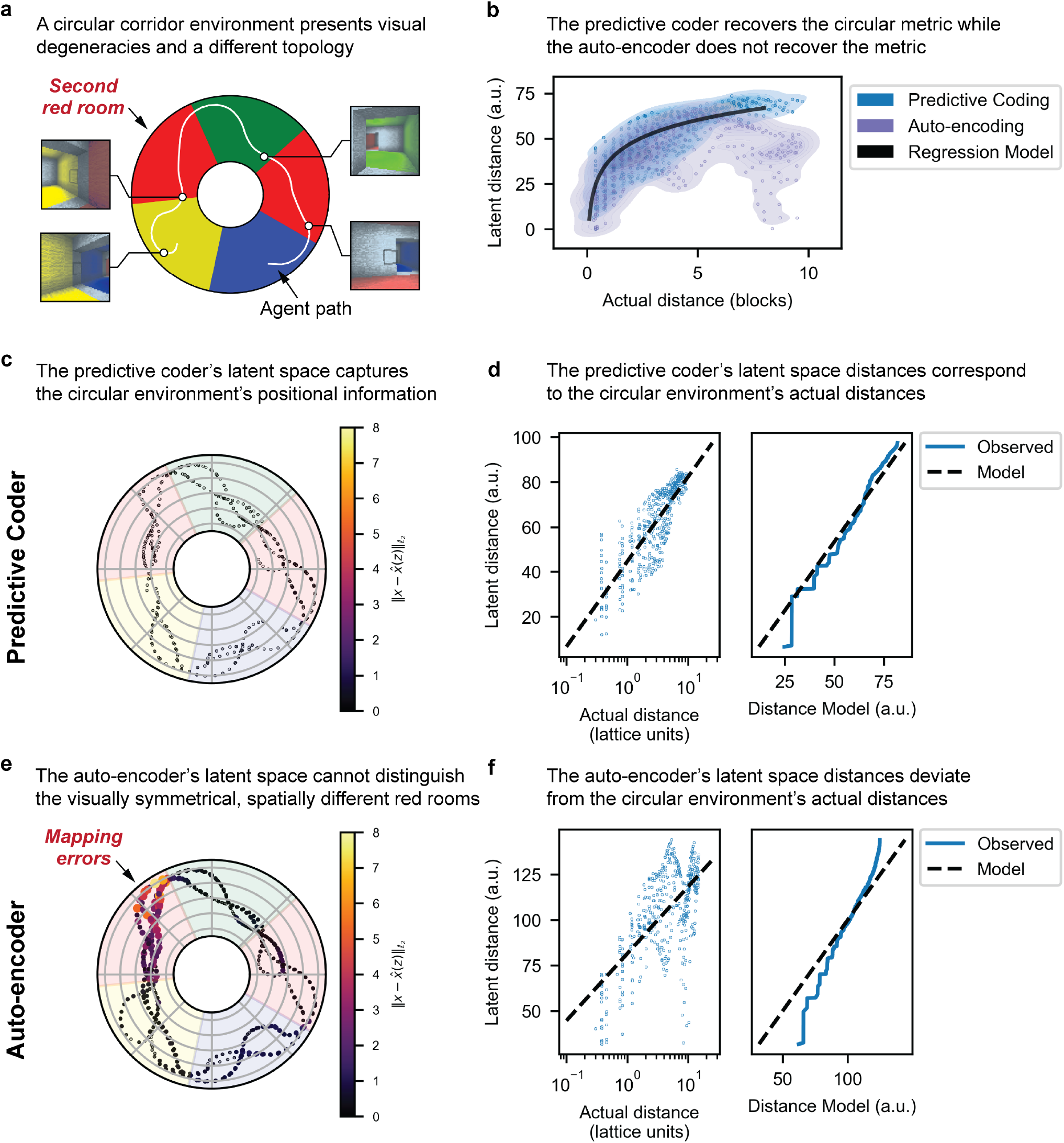
Predictive coding network can learn a circular topology and distinguishes visually identical, spatially different locations. **a**, an agent traverses a circular environment with two visually identical red rooms, which provides visually similar yet spatially different locations. **b**, the predictive coder’s latent distances show a correspondence with the circular environment’s metric while the auto-encoder’s latent distances show little correlation. **c**, similar to Figures 2 and 3, a different neural network measures the predictive coder’s spatial information by predicting the agent’s location from the predictive coder’s latent space. The predictive coder’s latent space demonstrates a low prediction error. **d**, similar to Figures 2 and 3, the nonlinear regression measures the correspondence between the latent distances ∥z − *z*^′^∥ and the actual distances ∥*x* − *x*^′^∥ with the model

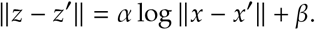

**d**, the correlation plot (left) with the nonlinear regression model show a strong correlation between the predictive coder’s latent distances and the environment’s actual distances (*r* = 0.827). The quantile-quantile plot (right) between the predictive coder’s latent distances and the regression model show high overlap (ⅅ _KL_ (*p*_PC_∥*p*_model_)= 0.250). **e**, without any past information, the auto-encoder cannot distinguish the two different red rooms and produces a high prediction error in these locations. **f**, the correlation plot (left) with the nonlinear regression model show little correlation between the auto-encoder’s latent distances and the environment’s actual distances (*r* = 0.288). The quantile-quantile plot (right) between the auto-encoder’s latent distances and the regression model show little overlap (ⅅ_KL_(*p*_PC_∥*p*_model_) = 3.806).

Similar to previous sections, we train a neural network (or a predictive coder) to perform predictive coding while traversing the circular corridor. In addition, we train a neural network (or an auto-encoder) to perform auto-encoding. The auto-encoder fails to recover spatial information in areas with visual degeneracy: it maps the two distinct red rooms to the same location (Figure 4(**e**)). In Figure 4(**e**), the auto-encoder predicts the images in the left red room to locations in the right red room—whereas locations with distinct visual scenes (such as the yellow and blue rooms) show a low prediction error (mean error 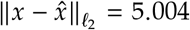 lattice units). In addition, the auto-encoder’s latent distances do not separate the different red rooms in latent space—whereas the predictive coder separates the two red rooms (Figure 4(**b**)). Moreover, the predictive coder demonstrates a low prediction error throughout, including the two visually degenerate red rooms (mean error 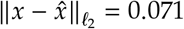 lattice units) (Figure 4(**c**)).

Moreover, we measure the relationship between the predictive coder’s (and auto-encoder’s) metric and the environment’s metric by fitting a regression model (Figure 4(**b**))

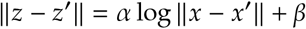

between the predictive coder’s (and auto-encoder’s) latent distances (∥z−*z*^′^∥) and the environment’s physical distances (*x*−*x*^′^). Compared to the natural environment, the auto-encoder’s latent distances shows more deviation from the environment’s spatial distances whereas the predictive coder’s latent distances maintain a correspondence with spatial distances. For the predictive coder, the latent metric recovers spatial metric quantitatively: the correlation plot (Figure 4(**d**, left)) shows a high correlation (*r* = 0.827) between the latent and spatial distances, and the quantile-quantile plot (Figure 4(**d**, right)) shows a high overlap between the regression model and the observed latent distances (ⅅ_KL_(*p*_PC_∥*p*_model_ = 0.250). The auto-encoder’s latent metric, conversely, does not recover the spatial metric: the correlation plot (Figure 4(**f**, left)) shows a low correlation (*r* = 0.288) between the latent and spatial distances, and the quantile-quantile plot (Figure 4(**f**, right)) shows a low overlap between the regression model and the observed latent distances (ⅅ_KL_(*p*_PC_∥*p*_model_) = 3.806).

As shown in Figure 4, the auto-encoder cannot recover the spatial map of the circular corridor—whereas the predictive coder can recover the map. Here we show that auto-encoders cannot the environment’s map for *any* environment with visual degeneracy, not just the circular corridor. To show that the auto-encoder cannot learn the environment’s map, we show that *any* statistical estimator cannot learn the environment’s map from *stationary* observations. For clarity and brevity, we will provide a proof sketch on a lattice environment *X*—a closed subset of ℤ ^2^.

#### Theorem 1.

*Consider an environment X—a closed subset of the lattice* ℤ ^2^ *with a function* 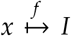 *that gives an image I*_*x*_ = *f*(*x*) ⊂ ℝ^*D*^ *for each position x*∈ *X. Let the environment’s observations be degenerate such that*

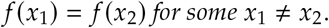

*There exists no decoder* 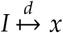 *that satisfies*

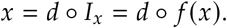

*Proof*. The proof proceeds as a consequence that a function has no left inverse *if and only if* it is not one-to-one.

Suppose there exists a decoder 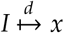 that satisfies

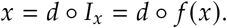

Consider

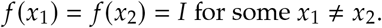

Then,

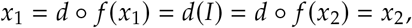

which is a contradiction, as required.

Because Theorem 1 demonstrates there exists no decoder for a visually degenerate environment with *stationary* observations, an auto-encoder *cannot* recover a visually degenerate environment; the auto-encoder’s failure arises because two locations with the same observation cannot be discriminated.

#### Corollary 1.

*Consider an auto-encoder g* = *dec* º *enc with an encoder* 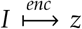 *and decoder* 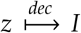 *that compresses images into a latent space z* ∈ ℝ^*H*^. *There exists no decoder* 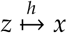 *that satisfies*

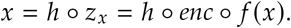

*Proof*. Consider the decoder *d* = *h* ºenc : *I* →*x*. By Theorem 1, this decoder cannot satisfy

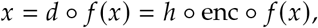

as required.

### Predictive coding generates units with localized receptive fields that support vector navigation

In the previous section, we demonstrate that the predictive coding neural network captures spatial relationships within an environment containing more internal spatial information than can be captured by an auto-encoder network that encodes image similarity. Here, we analyze the structure of the spatial code learned by the predictive coding network. We demonstrate that each unit in the neural network’s latent space activates at distinct, localized regions—akin to place fields in the mammalian brain—in the environment’s physical space (Figure 5(**a**)). These place fields overlap and their aggregate covers the entire physical space. Each physical location, is represented by a unique combination of overlapping regions encoded by the latent units. This combination of overlapping regions recovers the agent’s current physical position. Furthermore, given two physical locations, there now exist two distinct combinations of overlapping regions in latent space. Vector navigation is the representation of the vector heading to a goal location from a current location^51^. We show that overlapping regions (or place fields) can give a heading from a current locations to a goal location. Specifically, a linear decoder recovers the vector to a goal location from a starting location by taking the difference in place fields, which supports vector navigation^‡^ (Figure S1).

**Figure 5.**
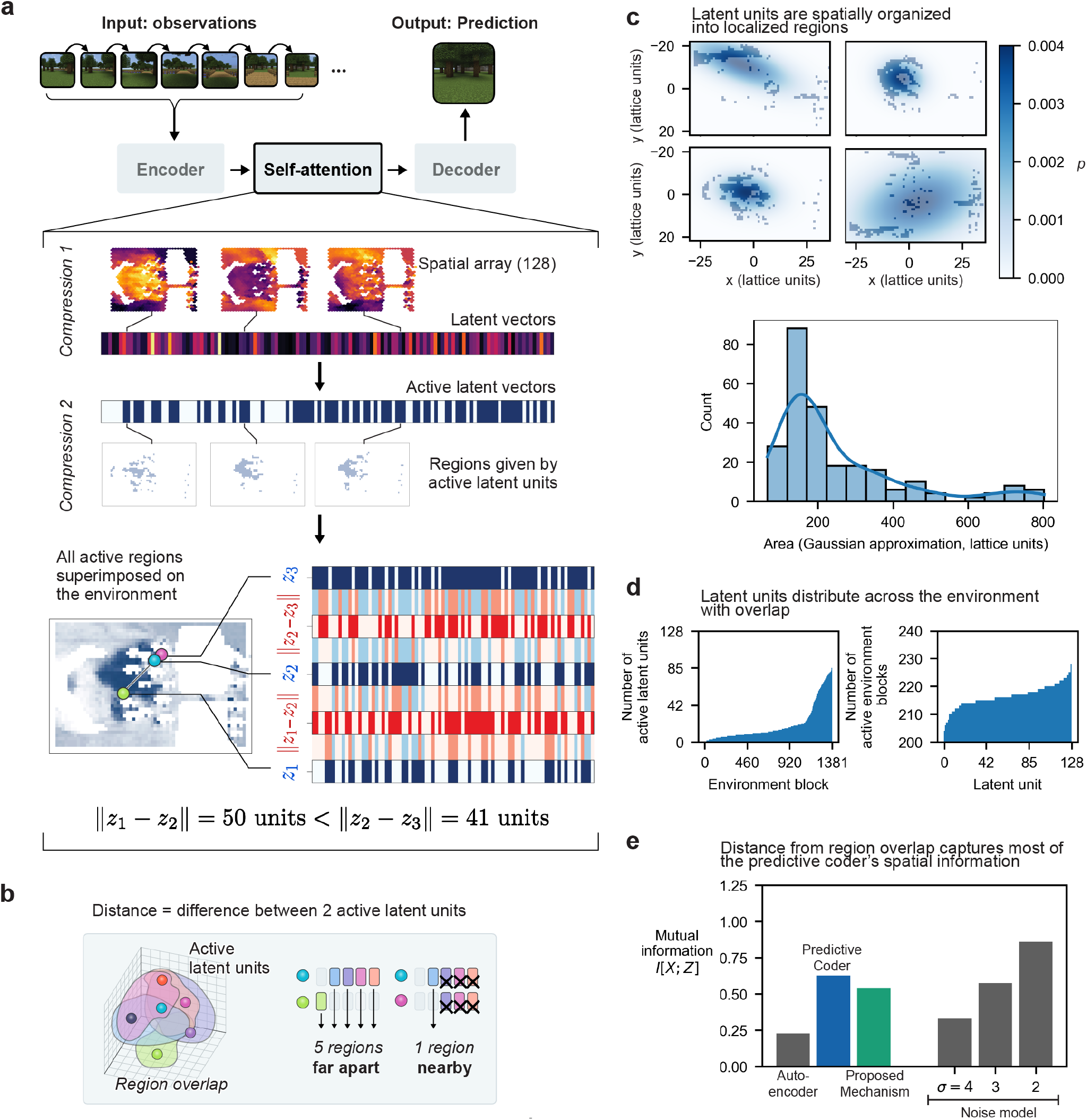
The predictive coding network generates place fields that support vector based distance calculations. **a**, when encoding past images for predictive coding, the self-attention module generates latent vectors. Each continuous unit in these latent vectors activates in concentrated, localized regions in physical space. These continuous units can be thresholded to generate a binary vector determining whether each unit is active. Each latent unit covers a unique region, and each physical location gives a unique combination of these overlapping regions. As an agent moves away from its original location, the combination of overlapping regions gradually deviates from its original combinations. This deviation, as measured by Hamming distance, correlates with physical distance. **b**, distance is given by the difference in the latent units’ overlapping regions. Two nearby locations have small deviations in overlap (right) while two distant locations have large deviations (middle). **c**, latent units are spatially organized into localized regions. The active latent units are approximated by a two-dimensional Gaussian distribution to measure the latent unit’s localization (top). The latent units’ Gaussian approximations are highly localized with a mean area of 254.6 for densities *p*≥0.0005. **d**, latent units distributed across the environment. The number of latent units was calculated as each lattice block in the environment (left), and the number of lattice blocks were calculated for each active unit (right). The latent units provide a unique combination for 87.6% of the environment, and their aggregate covers the entire environment. **e**, distance from the region overlap captures most of the predictive coder’s spatial information. We calculate the distance for every pair of active latent vectors and their respective physical Euclidean distances as a joint distribution. The proposed mechanism captures a majority of the predictive coder’s spatial information—as the proposed mechanism’s mutual information (0.542 bits) compares to the predictive coder’s mutual information (0.627 bits)

To support this proposed mechanism, we first demonstrate the neural network generates place fields. In other words, units from the neural network’s latent space produce localized regions in physical space. To determine whether a latent unit is active, we threshold the continuous value with its 90th-percentile value. The agent has head direction varies to ensure the regions are stable across all head directions. To measure a latent unit’s localization in physical space, we fit each latent unit distribution, with respect to physical space, to a two-dimensional Gaussian distribution (Figure 5(**c**), top)

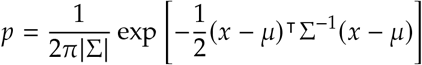

We measure the area of the ellipsoid given by the Gaussian approximation where *p* ≥ 0.0005 (Figure 5(**c**), bottom). The area of the latent unit approximation measures how localized a unit is compared to the environment’s area, which measures 40×65 = 2, 600 lattice units. The latent unit approximations have a mean area of 254.6 lattice units and a 80% of areas are < 352.6 lattice units, which cover 9.79% and 13.6% of the environment, respectively.

The units in the neural network’s latent space provide a unique combinatorial code for each spatial position. The aggregate of latent units covers the environment’s entire physical space. At each lattice block in the environment, we calculate the number of active latent units (Figure 5(**d**), left). The number of active latent units is different in 87.6% of the lattice blocks. Every lattice block has at least one active latent unit, which indicates the aggregate of the latent units cover the environment’s physical space. Moreover, to ensure the regions remain stable across shifting landmarks, environment’s trees were removed and randomly redistributed in the environment (Figure S5(**a-b**)). The regions remain stable after changing the tree landmarks with the Jaccard index (|*S*_new_∩*S*_old_|/|*S*_new_∪*S*_old_|) (or the intersection over union between new regions *S*_new_ and old *S*_old_ regions) of 0.828.

Lastly, we demonstrate that the neural network can measure physical distances and could perform vector navigation—representing the vector heading from a current location to a goal location—by comparing the combinations of overlapping regions in its latent space. We first determine the active latent units by thresholding each continuous value by its 90th-percentile value. At each position, we have a 128-dimensional binary vector that gives the overlap of 128 latent units. We take the bitwise difference *z*_1_ −*z*_2_ between the overlapping codes *z*_1_ and *z*_2_ at two varying positions *x*_1_ and *x*_2_ with the vector displacement *x*_1_ − *x*_2_ (Figure S1(**a**)). We then fit a linear decoder from the code *z*_1_ − *z*_2_ to the vector displacement *x*_1_ − *x*_2_,

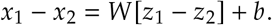

The predicted distance error 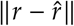 and the predicted direction error 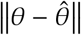 are decomposed from the predicted displacement 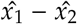. The linear decoder has a low prediction error for distance (< 80%, 12.49 lattice units; mean 7.89 lattice units) and direction (< 80%, 48.04°; mean 30.6°) (Figure S1(**b-c**)). The code *z*_1_ −*z*_2_ is highly correlated with direction *θ* and distance *r* with Pearson correlation coefficients 0.924 and 0.718, respectively (Figure S1(**d**)).

We can measure the correspondence between the bitwise distance^§^| *z*_1_ −*z*_2_ |and the physical distances ∥*x*_1_−*x*_2_ ∥ℓ_2_. Similar the previous sections, we compute the joint densities of the binary vectors’ bitwise distances and the physical positions’ Euclidean distances. We then calculate their mutual information to measure how much spatial information the bitwise distance captures. The proposed mechanism for the neural network’s distance measurement—the binary vector’s Hamming distance—gives a mutual information of 0.542 bits, compared to the predictive coder’s mutual information of 0.627 bits and the auto-encoder’s mutual information of 0.227 bits (Figure 5(**e**)). The code from the overlapping regions capture a majority amount of the predictive coder’s spatial information.

## Discussion

Mapping is a general mechanism for generating an internal representation of sensory information. While spatial maps facilitate navigation and planning within an environment, mapping is a ubiquitous neural function that extends to representations beyond visual-spatial mapping. The primary sensory cortex (S1), for example, maps tactile events topographically. Physical touches that occur in proximity are mapped in proximity for both the neural representations and the anatomical brain regions^52^. Similarly, the cortex maps natural speech by tiling regions with different words and their relationships, which shows that topographic maps in the brain extend to higher-order cognition. Similarly, the cortex maps natural speech by tiling regions with different words and their relationships, which shows that topographic maps in the brain extend to higher-order cognition. The similar representation of non-spatial and spatial maps in the brain suggests a common mechanism for charting cognitive^53^. However, it is unclear how a single mechanism can generate both spatial and non-spatial maps.

Here, we show that predictive coding provides a basic, general mechanism for charting spatial maps by predicting sensory data from past sensory experiences— including environments with degenerate observations. Our theoretical framework applies to any vector valued sensory data and could be extended to auditory data, tactile data, or tokenized representations of language. We demonstrate a neural network that performs predictive coding can construct an implicit spatial map of an environment by assembling information from local paths into a global frame within the neural network’s latent space. The implicit spatial map depends specifically on the sequential task of predicting future visual images. Neural networks trained as auto-encoders do not reconstruct a faithful geometric representation in the presence of physically distant yet visually similar landmarks.

Moreover, we study the predictive coding neural network’s representation in latent space. Each unit in the network’s latent space activates at distinct, localized regions—called place fields—with respect to physical space. At each physical location, there exists a unique combination of overlapping place fields. At two locations, the differences in the combinations of overlapping place fields provides the distance between the two physical locations. The existence of place fields in both the neural network and the hippocampus^16^ suggest that predictive coding is a universal mechanism for mapping. In addition, vector navigation emerges naturally from predictive coding by computing distances from overlapping place field units. Predictive coding may provide a model for understanding how place cells emerge, change, and function.

Predictive coding can be performed over any sensory modality that has some temporal sequence. As natural speech forms a cognitive map, predictive coding may underlie the geometry of human language. Intriguingly, large language models train on causal word prediction, a form of predictive coding, build internal maps that support generalized reasoning, answer questions, and mimic other forms of higher order reasoning^54^. Similarities in spatial and non-spatial maps in the brain suggest that large language models organize language into a cognitive map and chart concepts geometrically. These results all suggest that predictive coding might provide a unified theory for building representations of information—connecting disparate theories including place cell formation in the hippocampus, somatosensory maps in the cortex, and human language.

## Acknowledgements

We deeply appreciate Inna Strazhnik for her exceptional contributions to the scientific visualizations and figure illustrations. Her expertise in translating our research into clear visuals has significantly elevated the clarity and impact of our paper. We express our heartfelt gratitude to Thanos Siapas, Evgueniy Lubenov, Dean Mobbs, and Matthew Rosenberg for their invaluable and insightful discussions which profoundly enriched our work. Their expertise and feedback have been instrumental in the development and realization of this research. Additionally, we appreciate the insights provided by Lixiang Xu, Meng Wang, and Jieyu Zheng, which played a crucial role in refining various aspects of our study. The dedication and collaborative spirit of this collective group have truly elevated our research, and for that, we are deeply thankful.

## Code Availability

The code supporting the conclusions of this study is available on GitHub at https://github.com/jgornet/predictive-coding-recovers-maps. The repository contains the Malmo environment code, training scripts for both the predictive coding and autoencoding neural networks, as well as code for the analysis of predictive coding and autoencoding results. Should there be any questions or need for clarifications about the codebase, we encourage readers to raise an issue on the repository or reach out to the corresponding author.

## Data Availability

All datasets supporting the findings of this study, including the latent variables for the autoencoding and predictive coding neural networks, as well as the training and validation datasets, are available on GitHub at https://github.com/jgornet/predictive-coding-recovers-maps. Researchers and readers interested in accessing the data for replication, verification, or further studies can contact the corresponding author or refer to the supplementary materials section for more details.

## Notes

^1^Regarding the increasing number of channels in the middle, we initially ran experiments in the auto-encoder and the predictive coder with both the bottleneck architecture (decreasing middle channels) and the fan-out architecture (increasing middle channels). Both the bottleneck and fan-out architectures gave similar performance in visual prediction—for both the predictive coder and auto-encoder. In the predictive coder, we found that the latent variables in the fan-out architecture generate place cell-like fields seen in Figure 4. Because the auto-encoder has similar performance for the bottleneck and fan-out architectures, we use the fan-out architecture for the auto-encoder to provide a controlled comparison between auto-encoding and predictive coding.

^2^The appearance of place cell-like firing is common in simpler networks that perform spatial navigation such as Treves Miglino, Parisi (2007) and Sprekeler & Wiskott (2007). It is currently unclear whether artificial place cell-like behavior corresponds to biological place cells. Artificial place cell-like behavior could be an artifact of the simplified inputs or spatial coordinates.

## Supplementary Information

### Neural networks solve predictive coding by performing maximum likelihood estimation

We can express the model distribution *p*_*θ*_(*o*_*t*_ |*o*_<*t*_) as

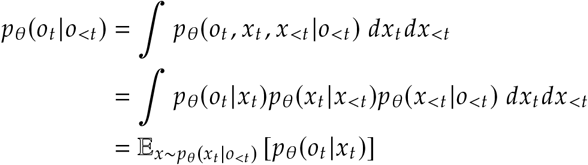

Performing maximum likelihood estimation,

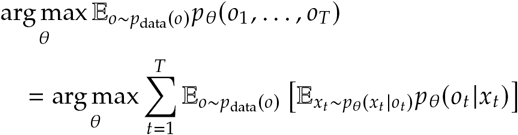

As log is a monotonic, increasing function, we can take perform maximum *log-likelihood* estimation,

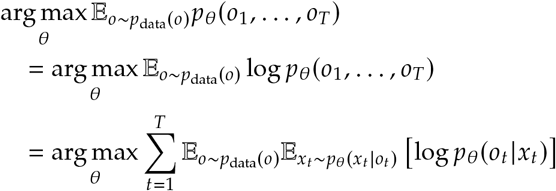

Predictive coding, which is solving for *p*_*θ*_(*o*_*t*_|*x*_*t*_) and *p*_*θ*_(*x*_*t*_ |*o*_<*t*_), is equivalent to estimating the data-generating distribution *p*_*θ*_(*o*_1_, …, *o*_*T*_).

Suppose that the agent’s path and observations are deterministic. First, the agent’s next position given its past positions

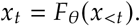

Second, the agent’s *sequence* of past positions given its *sequence* of observations is given by

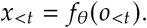

We can then parameterize the model distributions

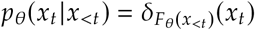

and

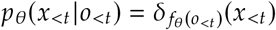

with neural networks 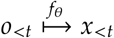 and 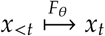. If every positions have a single observation, we can parameterize the distribution *p*_*θ*_(*o*_*t*_ |*x*_*t*_) as

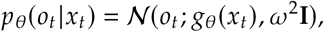

then maximum likelihood estimation becomes

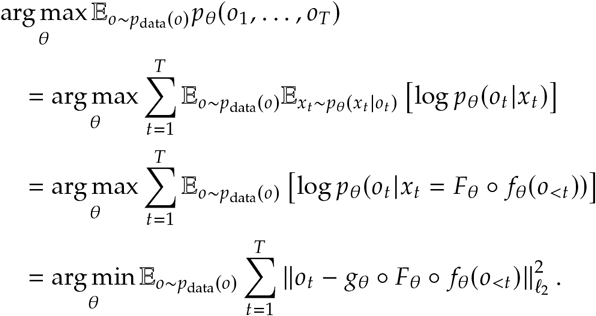

In neural networks, the errors are propagated by gradient descent

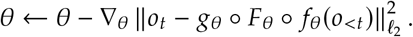

**Figure S1.**
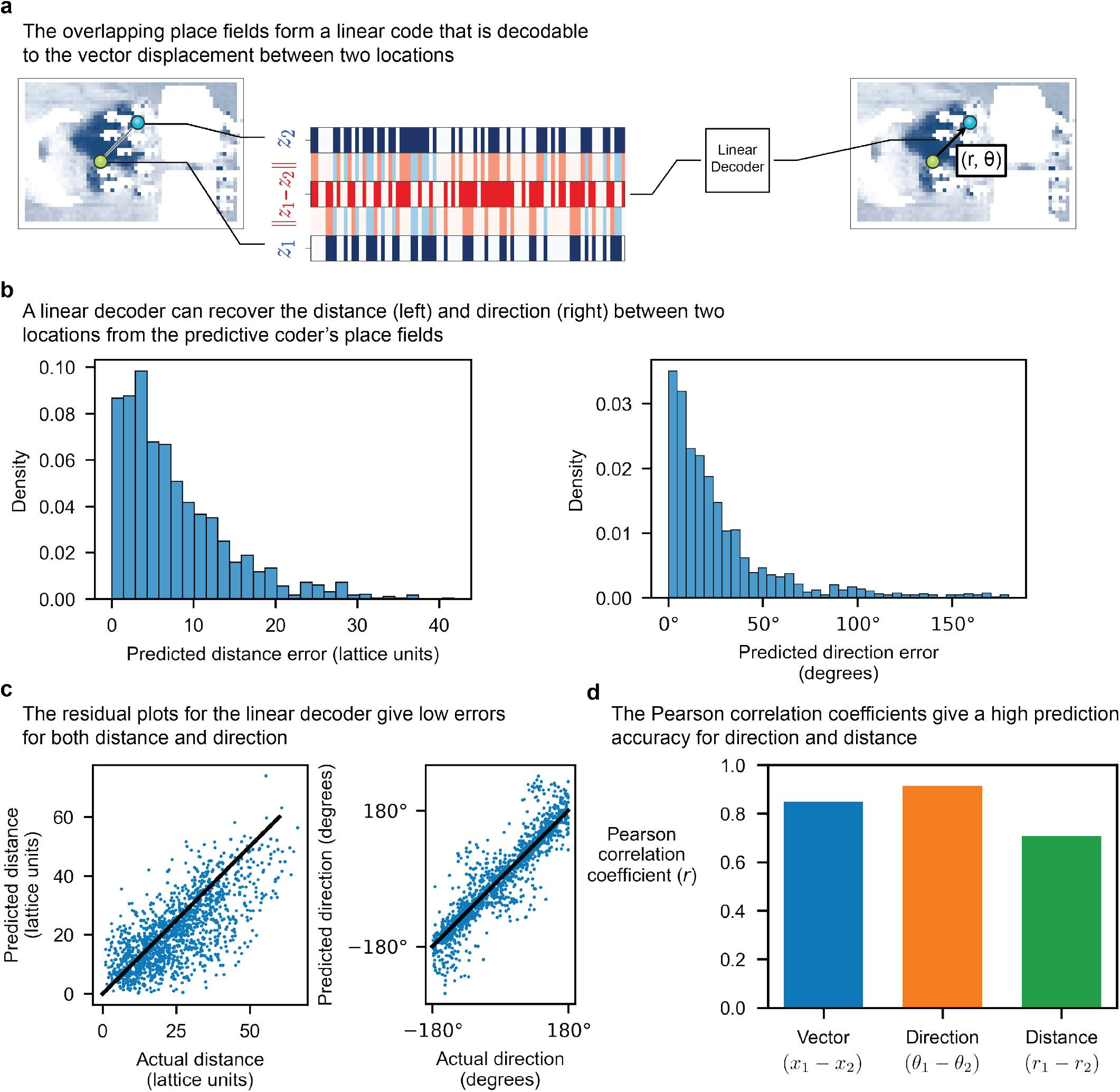
The place field overlap between two locations is linearly decodable to a vector heading. **a**, at different two locations *x*_1_ and *x*_2_, there exists a place field code *z*_1_ and *z*_2_, respectively. The bitwise different *z*_1_ − *z*_2_ gives the overlap between place fields at locations *x*_1_ and *x*_2_. We perform linear regression inputting the overlap codes *z*_1_−*z*_2_ and predicting the vector displacement *x*_1_−*x*_2_ between the two locations,

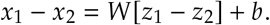

The linear model forms a linear decoder from the place field code *z*_1_ − *z*_2_ to the vector displacement *x*_1_ − *x*_2_. **b**, the errors in predicted distance 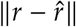 (left) and predicted direction 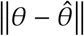 (right) are decomposed from the predicted displacement *x*_1_−*x*_2_. The linear decoder has a low prediction error for distance (< 80%, 12.49 lattice units; mean, 7.89 lattice units) and direction (< 80%, 48.04°; mean, 30.6°). **c**, residual plots show both distance (left) and direction (right) are strongly correlated with the place field code. **d**, in addition, the place field code is strongly correlated with the vector heading with Pearson correlation coefficients of 0.861, 0.718, and 0.924 for vector displacement (*x*_1_ −*x*_2_), distance (*r*_1_− *r*_2_), and direction (*θ*_1_− *θ*_2_), respectively.

**Figure S2.**
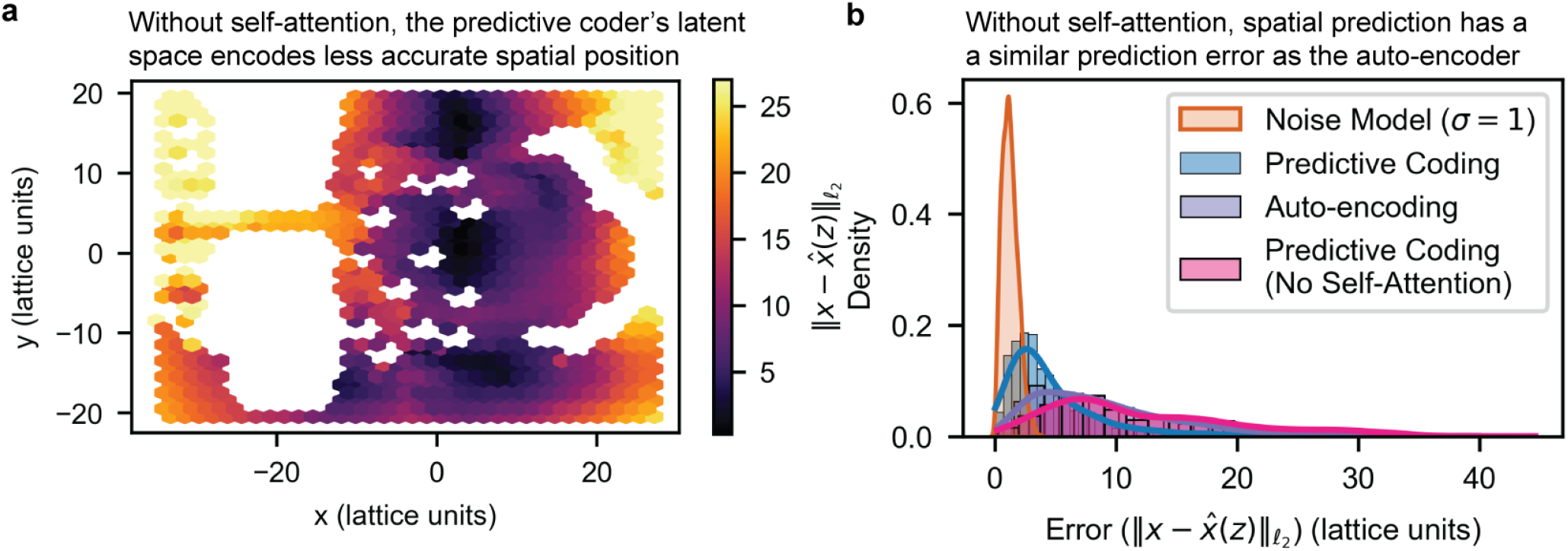
Without self-attention, the predictive coder encodes less accurate spatial information. **a-b**, self-attention in the predictive coder captures sequential information. To determine whether the temporal information is crucial to build an accurate spatial map, a neural network predicts the spatial location from the predictive coding’s latent space without self-attention. **a**, a heatmap of the predictive coder’s prediction error shows a low accuracy in many regions. **b**, the histogram of prediction errors shows a similar high prediction error as the auto-encoder.

### The predictive coding neural network requires self-attention for an accurate environmental map

The predictive coder architecture has three modules: encoder, self-attention, and decoder. The encoder and decoder act on single images—similar to how a tokenizer in language models transforms single words to vectors. The self-attention operates on image sequences to capture sequential patterns. If the predictive coder uses temporal structure to build a spatial map, then the self-attention should build the spatial map—not the encoder or decoder. In this section, we show that the temporal structure is required to build an accurate map of the environment.

Here we validate that self-attention is necessary to build an accurate map. First, we take the latent units encoded by the predictive coder’s encoder (without the self-attention). We then train a separate neural network to predict the actual spatial position given the latent unit. The accuracy of the predicted positions provides a lower bound on the spatial information given by the predictive coder’s encoder’s latent space. The heatmap (left) visualizes the errors given different positions in the environment. The histogram of the prediction errors (right) provides a comparison between the latent spaces of the predictive coder, the predictive coder without self–attention, and the auto-encoder. Without self-attention, the predictive coder has a much higher prediction error of its position similar to the auto-encoder.

**Figure S3.**
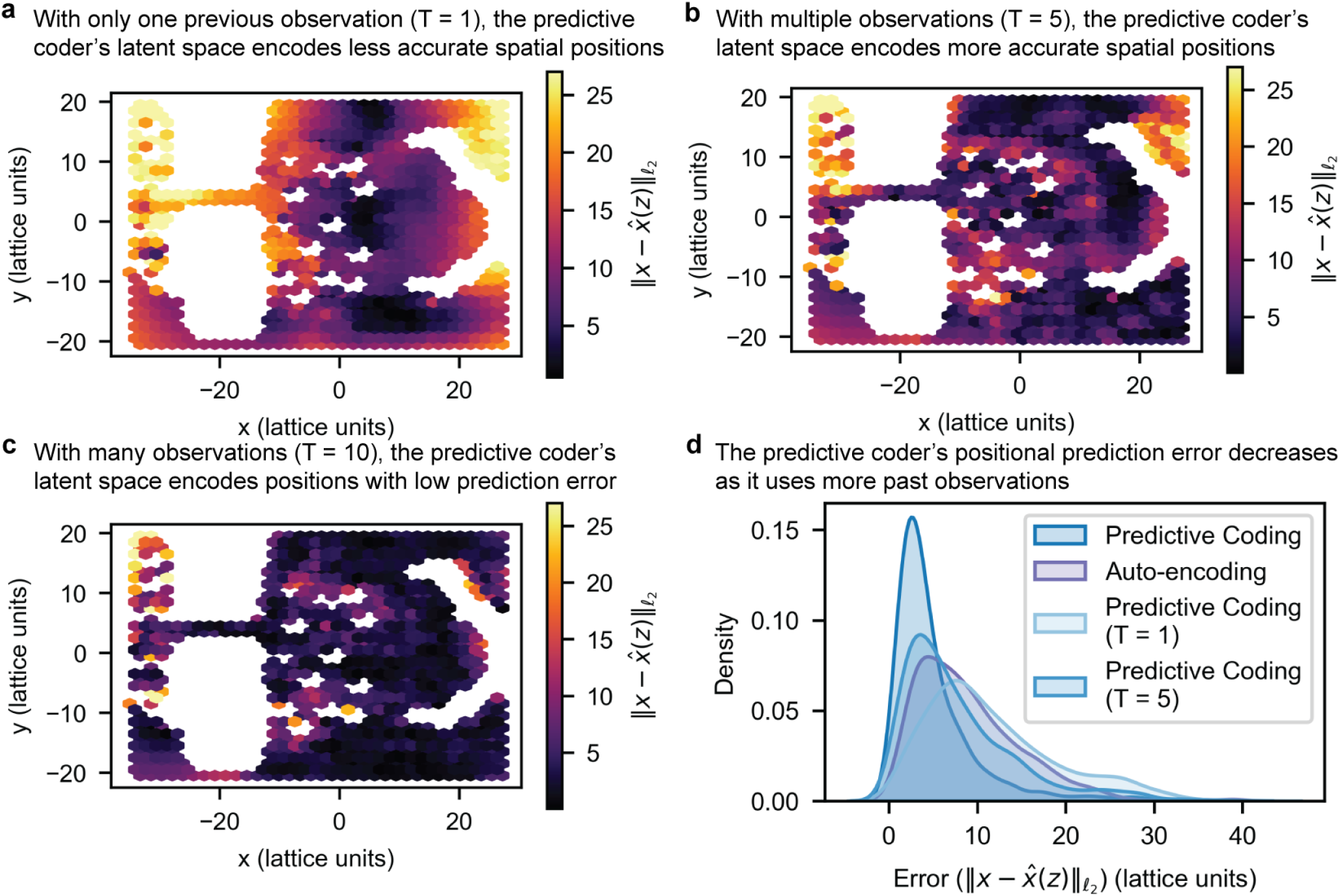
As the number of past observations increase, the predictive coder’s positional prediction error decreases. **a-c**, the predictive coder trains with one, five, and ten past observations, respectively. To determine how much temporal information is crucial to build an accurate spatial map, a neural network predicts the spatial location from the predictive coding’s latent space. **a**, with only one past observation, a heatmap of the predictive coder’s prediction error shows a high error in many regions. **b**, with five past observations, the prediction error reduces in many regions. **c**, with ten past observations, the prediction error is reduced below 7.3 lattice units for the majority (> 80%) of positions. **d**, as the number of past observations goes to zero, the histogram of prediction errors converges to the auto-encoder’s prediction error.

### The continuity and number of past observations determines the environmental map’s accuracy

In the predictive coder’s architecture, it contains three modules: the encoder, the self-attention, and the decoder. As discussed in the previous section, the predictive coder’s requires self-attention learn an accurate spatial map: the observation’s temporal information is crucial to build an environment’s map. A question that arises is how much temporal information does the predictive coder require to build an accurate map? In this section, we show that the predictive coder’s spatial prediction error decreases as the continuity and number of past observations.

First, we take the latent units encoded by the predictive coder’s encoder trained on differing numbers of past observations (*T* = 1, 5, 10). We then train a separate neural network to predict the actual spatial position given the latent unit. The accuracy of the predicted positions provides a lower bound on the spatial information given by the predictive coder’s encoder’s latent space. The heatmap (Figure S3(**a**, **b**, **c**)) visualizes the errors given different positions in the environment. With only one past observation, a heatmap of the predictive coder’s prediction error shows a high error in many regions. With five past observations, the prediction error reduces in many regions. With ten past observations, the prediction error is reduced below 7.3 lattice units for the majority (> 80%) of positions. As the number of past observations goes to zero, the histogram of prediction errors converges to the auto-encoder’s prediction error (Figure S3(**d**)).

**Figure S4.**
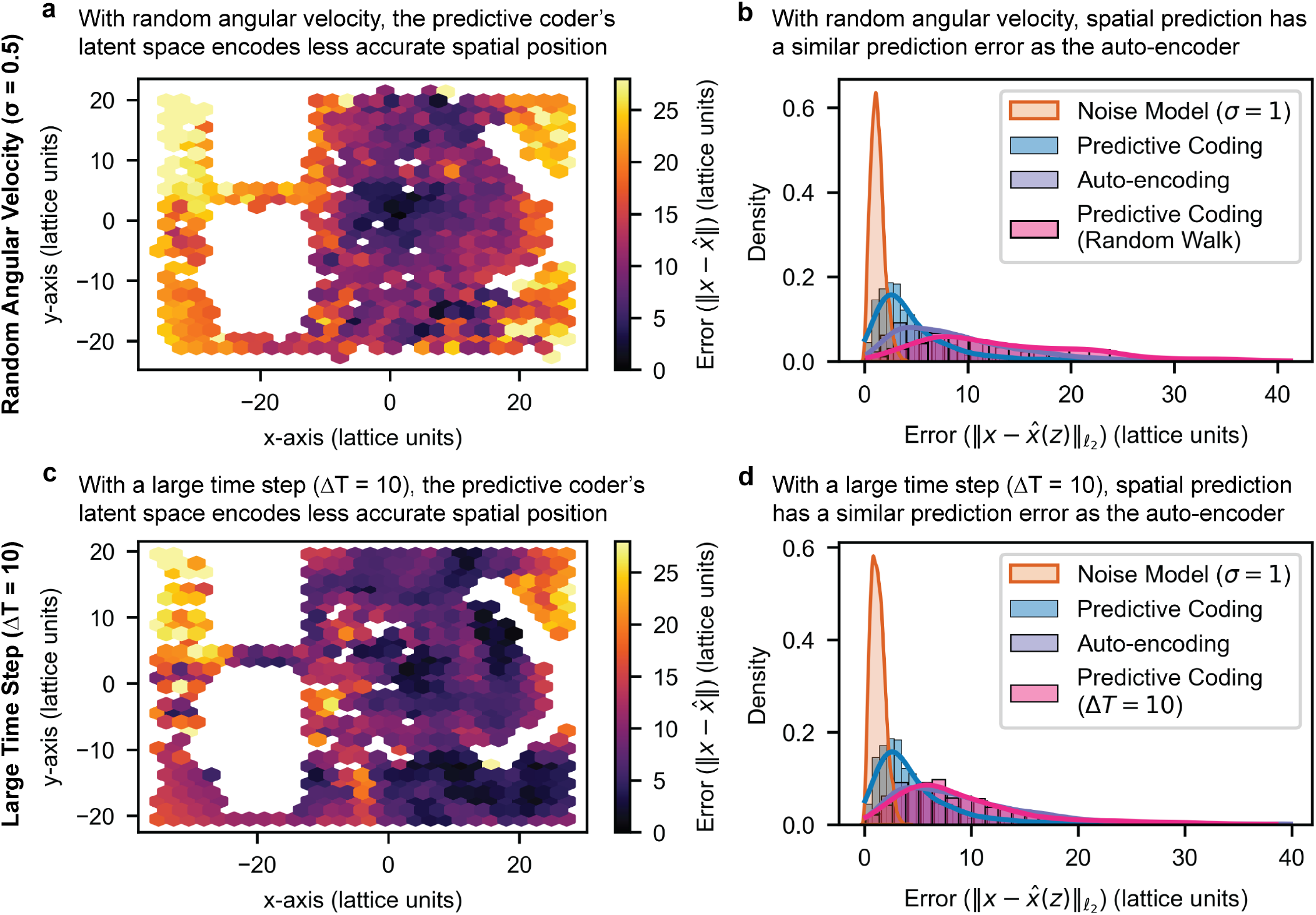
As the number of past observations increase, the predictive coder’s positional prediction error decreases (cont.). **a-b**, the agent typically traverses the environment in direct path with minimal angular rotation. To determine the impact of angular rotation on the predictive coder, the agent samples a random angular velocity (σ = ^30°^/_sec_) as it traverses the environment. The positional prediction error (**a**) increases and the error density **(b)** shifts to the auto-encoder’s error density. **c-d**, the agent typically takes short time steps per an observation (20 images per second). To determine the impact of the time step length, the agent samples the environment’s images with a lower frame rate (2 images per second). Similar to the random angular velocity, the large time step results in the predictive coder having a higher prediction error **(c)** and an error density **(d)** shifting toward the auto-encoder’s error density.

**Figure S5.**
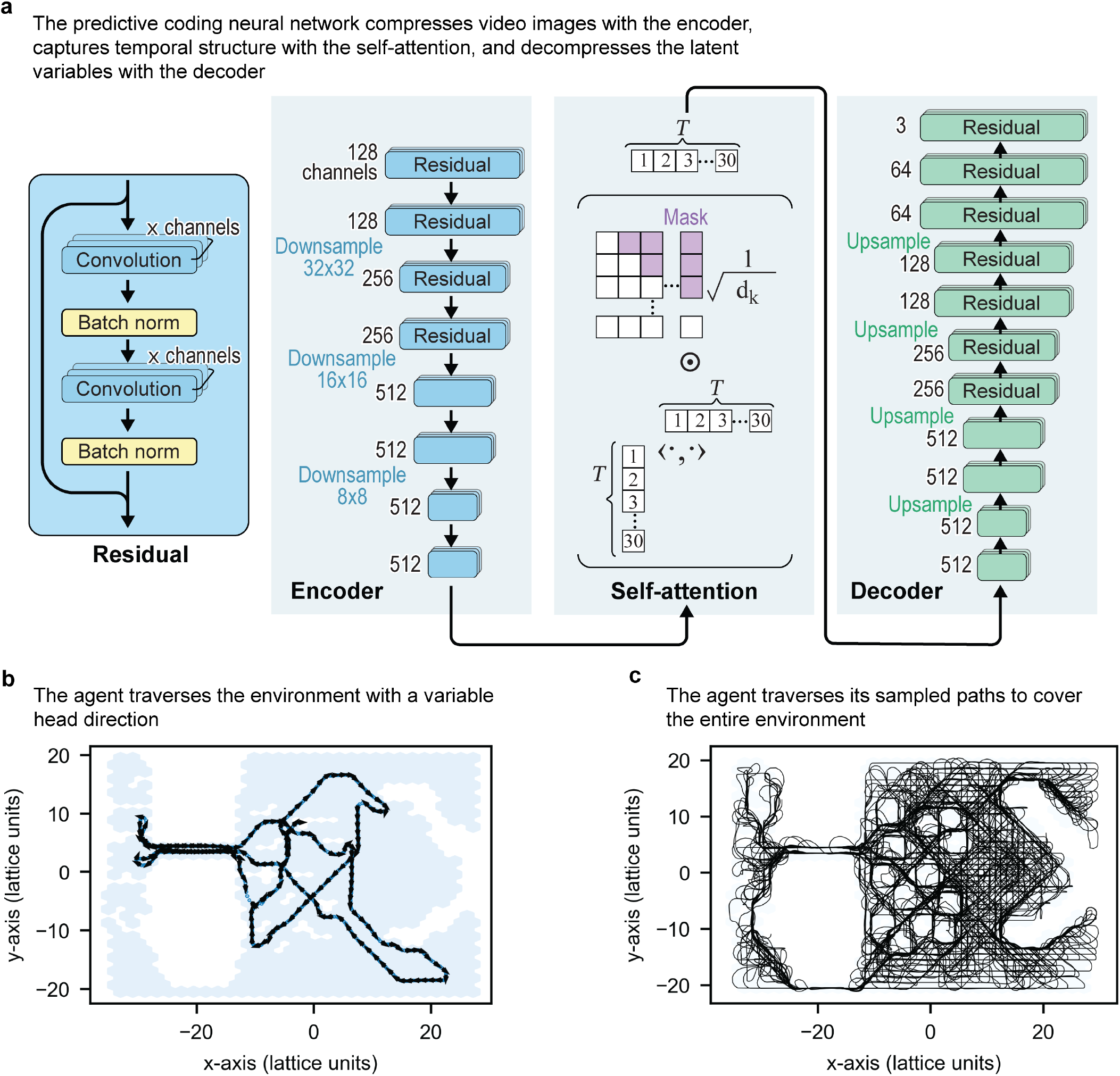
Extended neural network architecture and training description. **a**, the predictive coding neural network, or predictive coder, uses an encoder, self-attention, and decoder to perform predictive coding. The encoder is a convolutional neural network architecture called ResNet-18 that uses residuals to compress the high-dimensional video image. The self-attention module capture temporal dependencies from the low-dimensional encoded images. The self-attention’s output gives the predictive coder’s latent variables. The decoder is a convolutional neural network that upscales—rather than downscales—the predictive coder’s latent variables to predicted images. **b**, an example path of the agent shows the agent traversing the environment with a variable head direction. **c**, the agent’s paths traverse the space to cover the entire environment.

**Figure S6.**
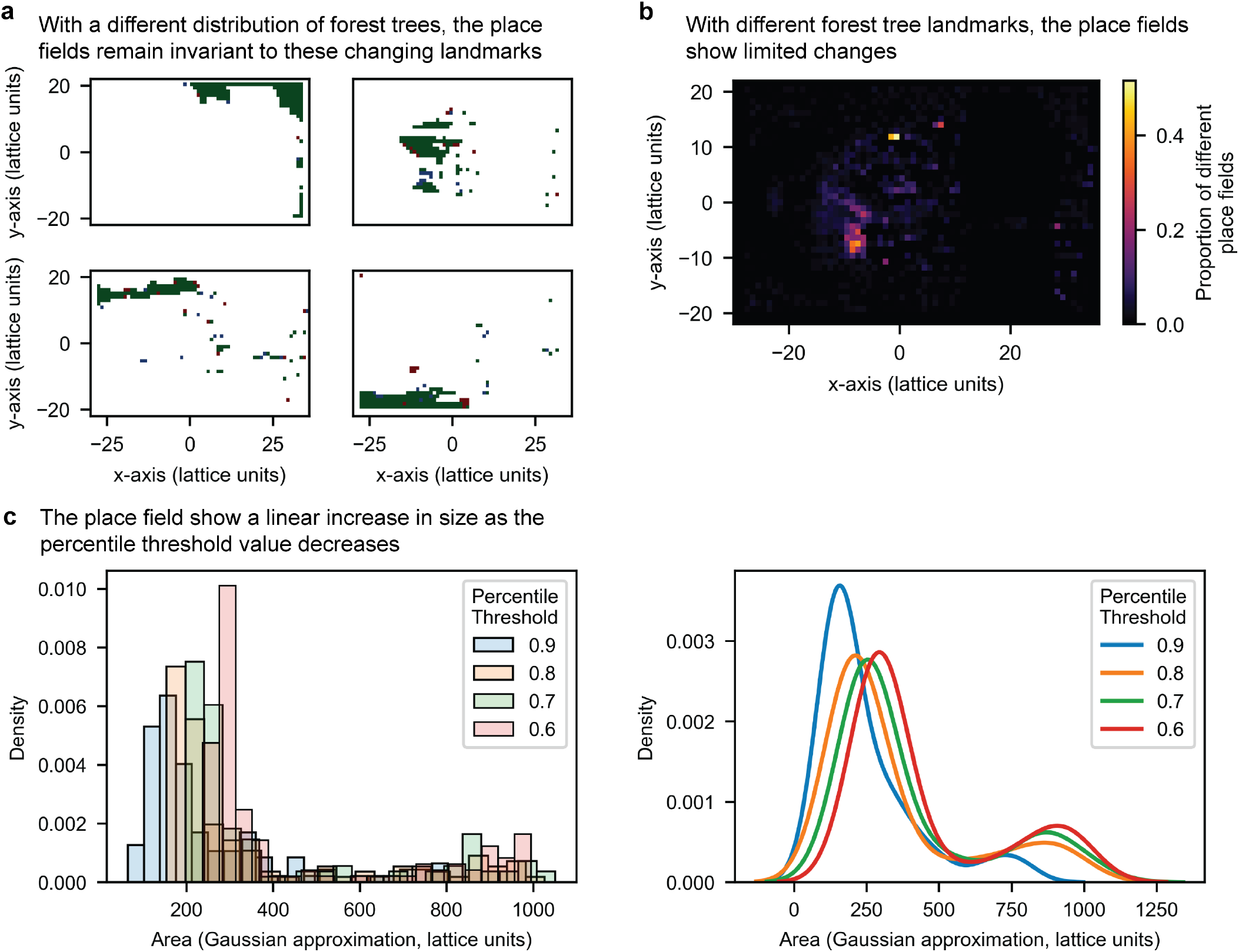
Extended place field results. **a-b**, The predictive coder’s place field latent variables are invariant to shifting landmarks. To determine the effect of shifting landmarks, the trees in the environment were removed and randomly redistributed in the forest region. **a**, the predictive coder’s original place fields (**red**) were overlaid with the new place fields (**blue**), and the union set of the original and new place fields are shown in green. The new, shifted landmark place fields demonstrate a large overlap (Jaccard index (|*A*∩*B*|/| *A* ∪*B*|) = 0.828) with the original place fields. **b**, a visual overlap of the proportion of different place fields at every location. The place fields show no variability outside the forest and low variability inside the forest region. **c**, the place fields are measured by thresholding the predictive coder’s latent variable, so the place field sizes are dependent on the percentile threshold value. To determine the effect of the threshold value on the place field sizes, the place field size histogram (left) is plotted with respect to the percentile threshold value, and the place field size densities (right) are estimated using kernel density estimation from the histogram.

### Footnotes

† The neural network is a feedforward deep neural network trained using backpropagation, or gradient descent, rather than Helmholtz machines^41,42^, which are commonly used in predictive coding.

‡ Studies such as Bush *et al*. (2015) consider grid cell-supported vector navigation whereas we only consider vector navigation using place cells.

§ In previous sections, we compute the Euclidean distance between latent units. For the bitwise distance, we threshold the latent units to its 90th-percentile then compute the c_1_-norm between the units.

