## Supplemental Information for "Automated construction of cognitive maps with visual predictive coding"

### Supplementary Information

#### Neural networks solve predictive coding by performing maximum likelihood estimation

We can express the model distribution $p_\theta(o_t|o_{<t})$ as

$$\begin{aligned} p_\theta(o_t|o_{<t}) &= \int p_\theta(o_t, x_t, x_{<t}|o_{<t})\ dx_t dx_{<t} \\ &= \int p_\theta(o_t|x_t)p_\theta(x_t|x_{<t})p_\theta(x_{<t}|o_{<t})\ dx_t dx_{<t} \\ &= \mathbb{E}_{x\sim p_\theta(x_t|o_{<t})}\left[p_\theta(o_t|x_t)\right] \end{aligned}$$

Performing maximum likelihood estimation,

$$\begin{aligned} &\arg\max_\theta \mathbb{E}_{o\sim p_{\text{data}}(o)} p_\theta(o_1, \ldots, o_T) \\ &= \arg\max_\theta \sum_{t=1}^{T} \mathbb{E}_{o\sim p_{\text{data}}(o)} \left[\mathbb{E}_{x_t\sim p_\theta(x_t|o_t)} p_\theta(o_t|x_t)\right] \end{aligned}$$

As log is a monotonic, increasing function, we can take perform maximum *log-likelihood* estimation,

$$\begin{aligned} &\arg\max_\theta \mathbb{E}_{o\sim p_{\text{data}}(o)} p_\theta(o_1, \ldots, o_T) \\ &= \arg\max_\theta \mathbb{E}_{o\sim p_{\text{data}}(o)} \log p_\theta(o_1, \ldots, o_T) \\ &= \arg\max_\theta \sum_{t=1}^{T} \mathbb{E}_{o\sim p_{\text{data}}(o)} \mathbb{E}_{x_t\sim p_\theta(x_t|o_t)} \left[\log p_\theta(o_t|x_t)\right] \end{aligned}$$

Predictive coding, which is solving for $p_\theta(o_t|x_t)$ and $p_\theta(x_t|o_{<t})$, is equivalent to estimating the data-generating distribution $p_\theta(o_1, \ldots, o_T)$.

Suppose that the agent's path and observations are deterministic. First, the agent's next position given its past positions

$$x_t = F_\theta(x_{<t}).$$

Second, the agent's *sequence* of past positions given its *sequence* of observations is given by

$$x_{<t} = f_\theta(o_{<t}).$$

We can then parameterize the model distributions

$$p_\theta(x_t|x_{<t}) = \delta_{F_\theta(x_{<t})}(x_t)$$

and

$$p_\theta(x_{<t}|o_{<t}) = \delta_{f_\theta(o_{<t})}(x_{<t})$$

with neural networks $o_{<t} \overset{f_\theta}{\longmapsto} x_{<t}$ and $x_{<t} \overset{F_\theta}{\longmapsto} x_t$. If every positions have a single observation, we can parameterize the distribution $p_\theta(o_t|x_t)$ as

$$p_\theta(o_t|x_t) = \mathcal{N}(o_t; g_\theta(x_t), \omega^2 \mathbf{I}),$$

then maximum likelihood estimation becomes

$$\begin{aligned} &\arg\max_\theta \mathbb{E}_{o\sim p_{\text{data}}(o)} p_\theta(o_1, \ldots, o_T) \\ &= \arg\max_\theta \sum_{t=1}^{T} \mathbb{E}_{o\sim p_{\text{data}}(o)} \mathbb{E}_{x_t\sim p_\theta(x_t|o_t)} \left[\log p_\theta(o_t|x_t)\right] \\ &= \arg\max_\theta \sum_{t=1}^{T} \mathbb{E}_{o\sim p_{\text{data}}(o)} \left[\log p_\theta(o_t|x_t = F_\theta \circ f_\theta(o_{<t}))\right] \\ &= \arg\min_\theta \mathbb{E}_{o\sim p_{\text{data}}(o)} \sum_{t=1}^{T} \|o_t - g_\theta \circ F_\theta \circ f_\theta(o_{<t})\|_{\ell_2}^2 . \end{aligned}$$

In neural networks, the errors are propagated by gradient descent

$$\theta \leftarrow \theta - \nabla_\theta \|o_t - g_\theta \circ F_\theta \circ f_\theta(o_{<t})\|_{\ell_2}^2 .$$

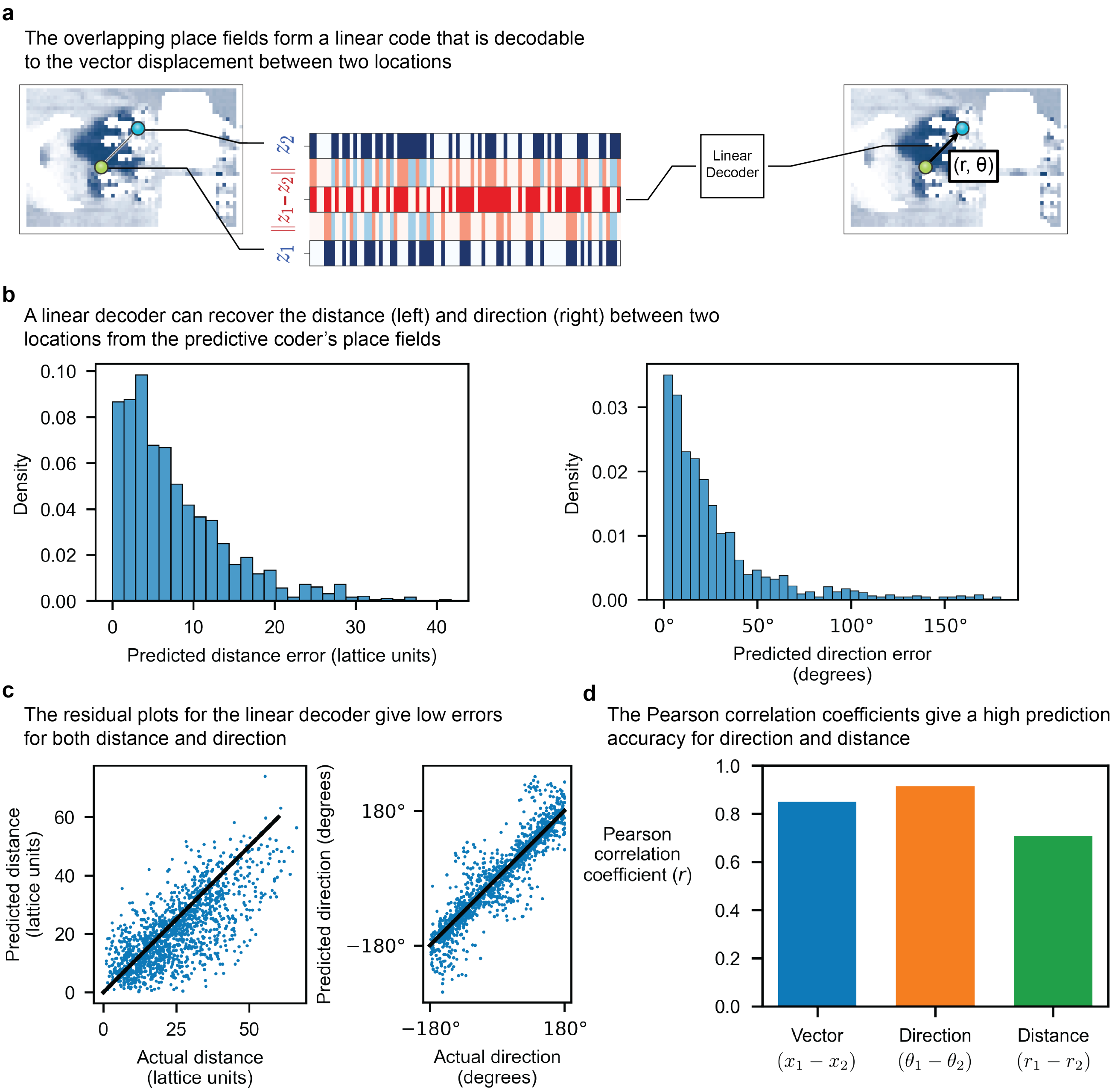


**Figure S1. The place field overlap between two locations is linearly decodable to a vector heading.** **a,** at different two locations $x_1$ and $x_2$, there exists a place field code $z_1$ and $z_2$, respectively. The bitwise different $z_1 - z_2$ gives the overlap between place fields at locations $x_1$ and $x_2$. We perform linear regression inputting the overlap codes $z_1 - z_2$ and predicting the vector displacement $x_1 - x_2$ between the two locations,

$$x_1 - x_2 = W[z_1 - z_2] + b.$$

The linear model forms a linear decoder from the place field code $z_1 - z_2$ to the vector displacement $x_1 - x_2$. **b,** the errors in predicted distance $\|r - \hat{r}\|$ (left) and predicted direction $\|\theta - \hat{\theta}\|$ (right) are decomposed from the predicted displacement $x_1 - x_2$. The linear decoder has a low prediction error for distance ($< 80\%$, 12.49 lattice units; mean, 7.89 lattice units) and direction ($< 80\%$, 48.04°; mean, 30.6°). **c,** residual plots show both distance (left) and direction (right) are strongly correlated with the place field code. **d,** in addition, the place field code is strongly correlated with the vector heading with Pearson correlation coefficients of 0.861, 0.718, and 0.924 for vector displacement ($x_1 - x_2$), distance ($r_1 - r_2$), and direction ($\theta_1 - \theta_2$), respectively.

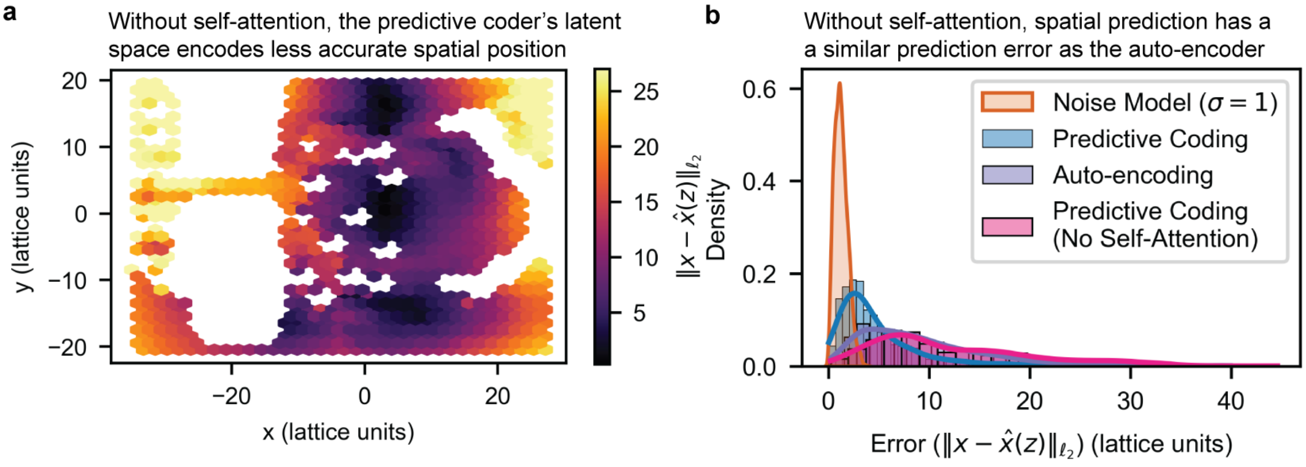


**Figure S2. Without self-attention, the predictive coder encodes less accurate spatial information.** **a-b,** self-attention in the predictive coder captures sequential information. To determine whether the temporal information is crucial to build an accurate spatial map, a neural network predicts the spatial location from the predictive coding's latent space without self-attention. **a,** a heatmap of the predictive coder's prediction error shows a low accuracy in many regions. **b,** the histogram of prediction errors shows a similar high prediction error as the auto-encoder.

---

#### The predictive coding neural network requires self-attention for an accurate environmental map

The predictive coder architecture has three modules: encoder, self-attention, and decoder. The encoder and decoder act on single images—similar to how a tokenizer in language models transforms single words to vectors. The self-attention operates on image sequences to capture sequential patterns. If the predictive coder uses temporal structure to build a spatial map, then the self-attention should build the spatial map—not the encoder or decoder. In this section, we show that the temporal structure is required to build an accurate map of the environment.

Here we validate that self-attention is necessary to build an accurate map. First, we take the latent units encoded by the predictive coder's encoder (without the self-attention). We then train a separate neural network to predict the actual spatial position given the latent unit. The accuracy of the predicted positions provides a lower bound on the spatial information given by the predictive coder's encoder's latent space. The heatmap (left) visualizes the errors given different positions in the environment. The histogram of the prediction errors (right) provides a comparison between the latent spaces of the predictive coder, the predictive coder without self–attention, and the auto-encoder. Without self-attention, the predictive coder has a much higher prediction error of its position similar to the auto-encoder.

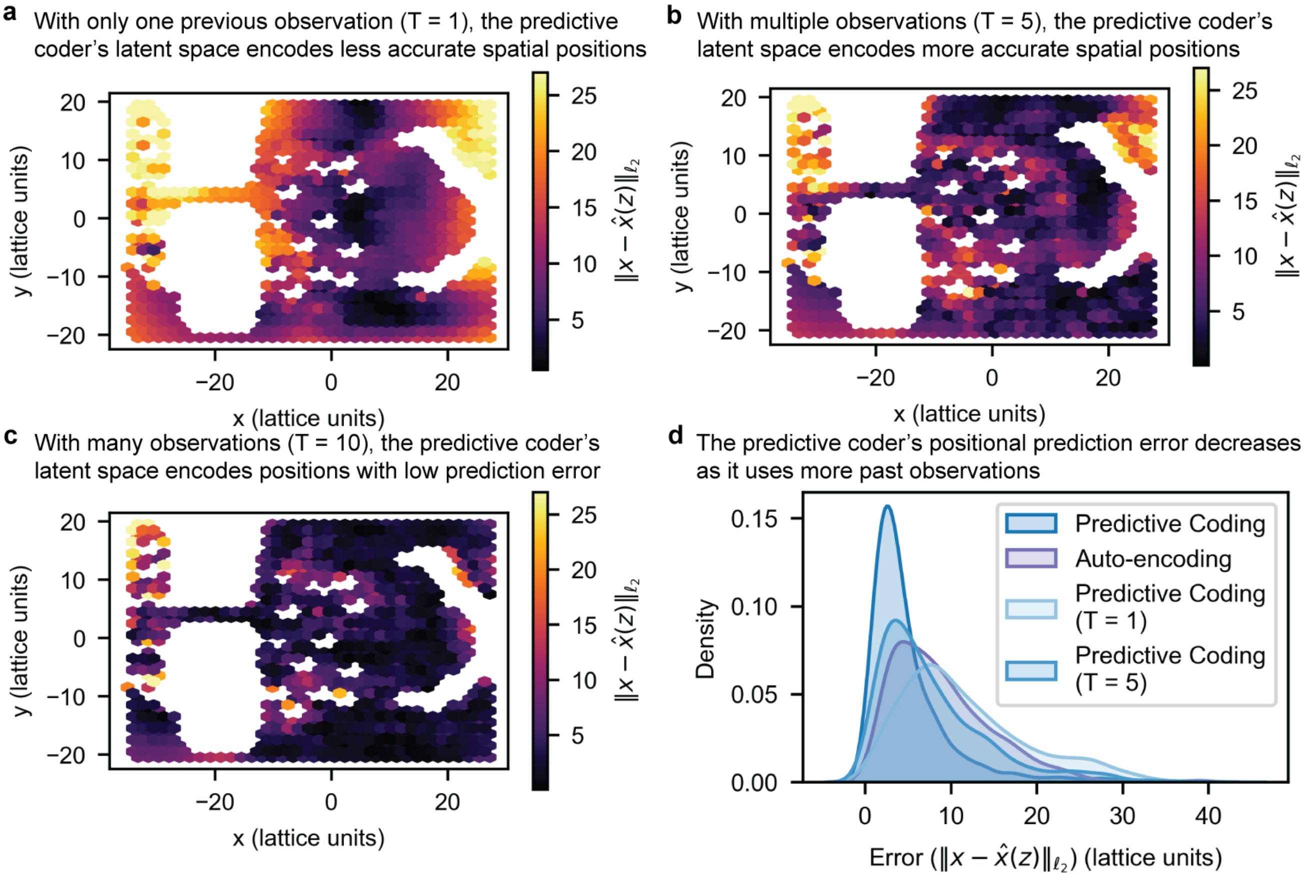


**Figure S3. As the number of past observations increase, the predictive coder's positional prediction error decreases.** **a-c,** the predictive coder trains with one, five, and ten past observations, respectively. To determine how much temporal information is crucial to build an accurate spatial map, a neural network predicts the spatial location from the predictive coding's latent space. **a,** with only one past observation, a heatmap of the predictive coder's prediction error shows a high error in many regions. **b,** with five past observations, the prediction error reduces in many regions. **c,** with ten past observations, the prediction error is reduced below 7.3 lattice units for the majority (> 80%) of positions. **d,** as the number of past observations goes to zero, the histogram of prediction errors converges to the auto-encoder's prediction error.

---

#### The continuity and number of past observations determines the environmental map's accuracy

In the predictive coder's architecture, it contains three modules: the encoder, the self-attention, and the decoder. As discussed in the previous section, the predictive coder's requires self-attention learn an accurate spatial map: the observation's temporal information is crucial to build an environment's map. A question that arises is how much temporal information does the predictive coder require to build an accurate map? In this section, we show that the predictive coder's spatial prediction error decreases as the continuity and number of past observations.

First, we take the latent units encoded by the predictive coder's encoder trained on differing numbers of past observations ($T = 1, 5, 10$). We then train a separate neural network to predict the actual spatial position given the latent unit. The accuracy of the predicted positions provides a lower bound on the spatial information given by the predictive coder's encoder's latent space. The heatmap (Figure S3(**a**, **b**, **c**)) visualizes the errors given different positions in the environment. With only one past observation, a heatmap of the predictive coder's prediction error shows a high error in many regions. With five past observations, the prediction error reduces in many regions. With ten past observations, the prediction error is reduced below 7.3 lattice units for the majority (> 80%) of positions. As the number of past observations goes to zero, the histogram of prediction errors converges to the auto-encoder's prediction error (Figure S3(**d**)).

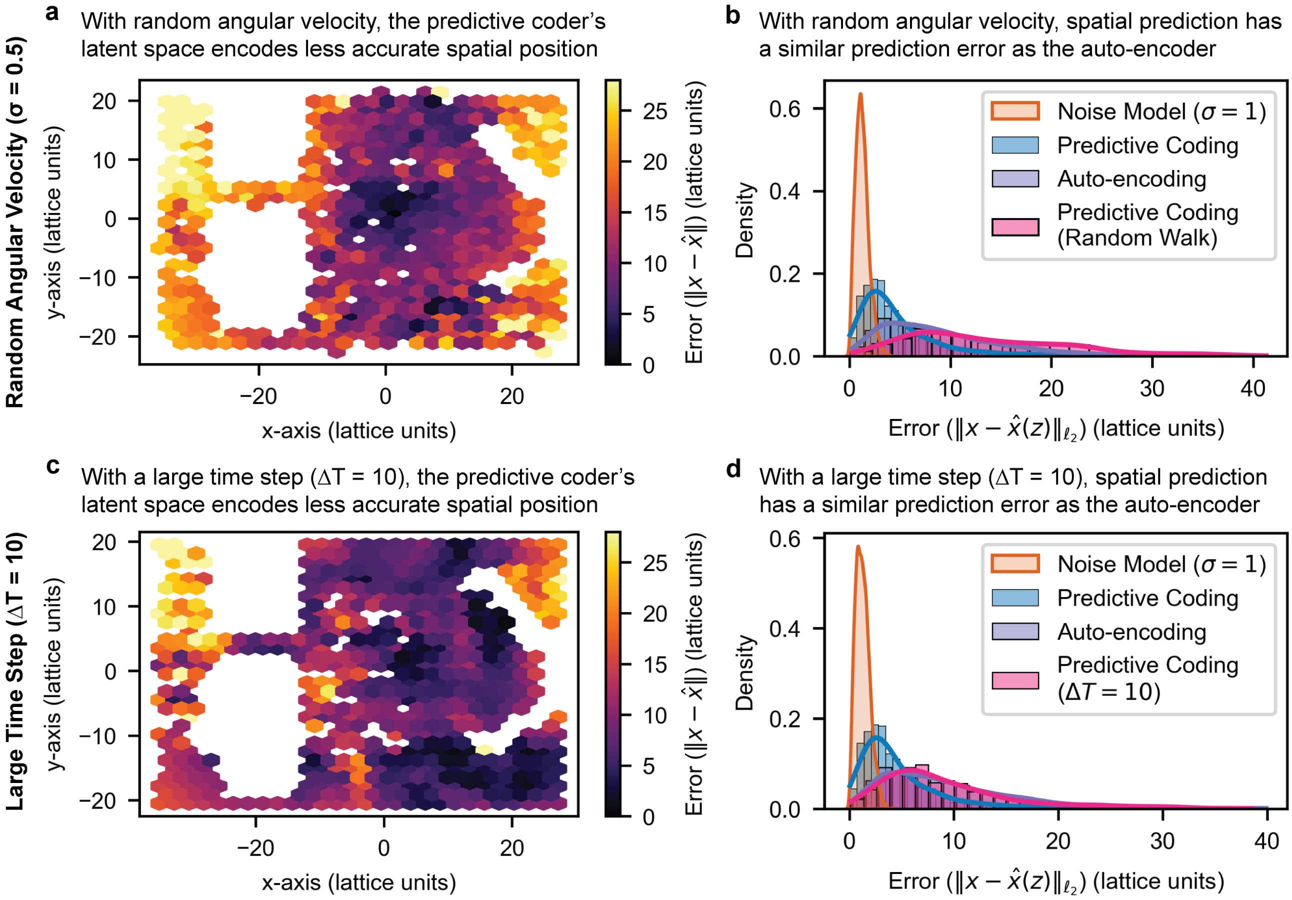


**Figure S4. As the number of past observations increase, the predictive coder's positional prediction error decreases (cont.). a-b,** the agent typically traverses the environment in direct path with minimal angular rotation. To determine the impact of angular rotation on the predictive coder, the agent samples a random angular velocity ($\sigma = {}^{30°}/_{\text{sec}}$) as it traverses the environment. The positional prediction error (**a**) increases and the error density **(b)** shifts to the auto-encoder's error density. **c-d,** the agent typically takes short time steps per an observation (20 images per second). To determine the impact of the time step length, the agent samples the environment's images with a lower frame rate (2 images per second). Similar to the random angular velocity, the large time step results in the predictive coder having a higher prediction error **(c)** and an error density **(d)** shifting toward the auto-encoder's error density.

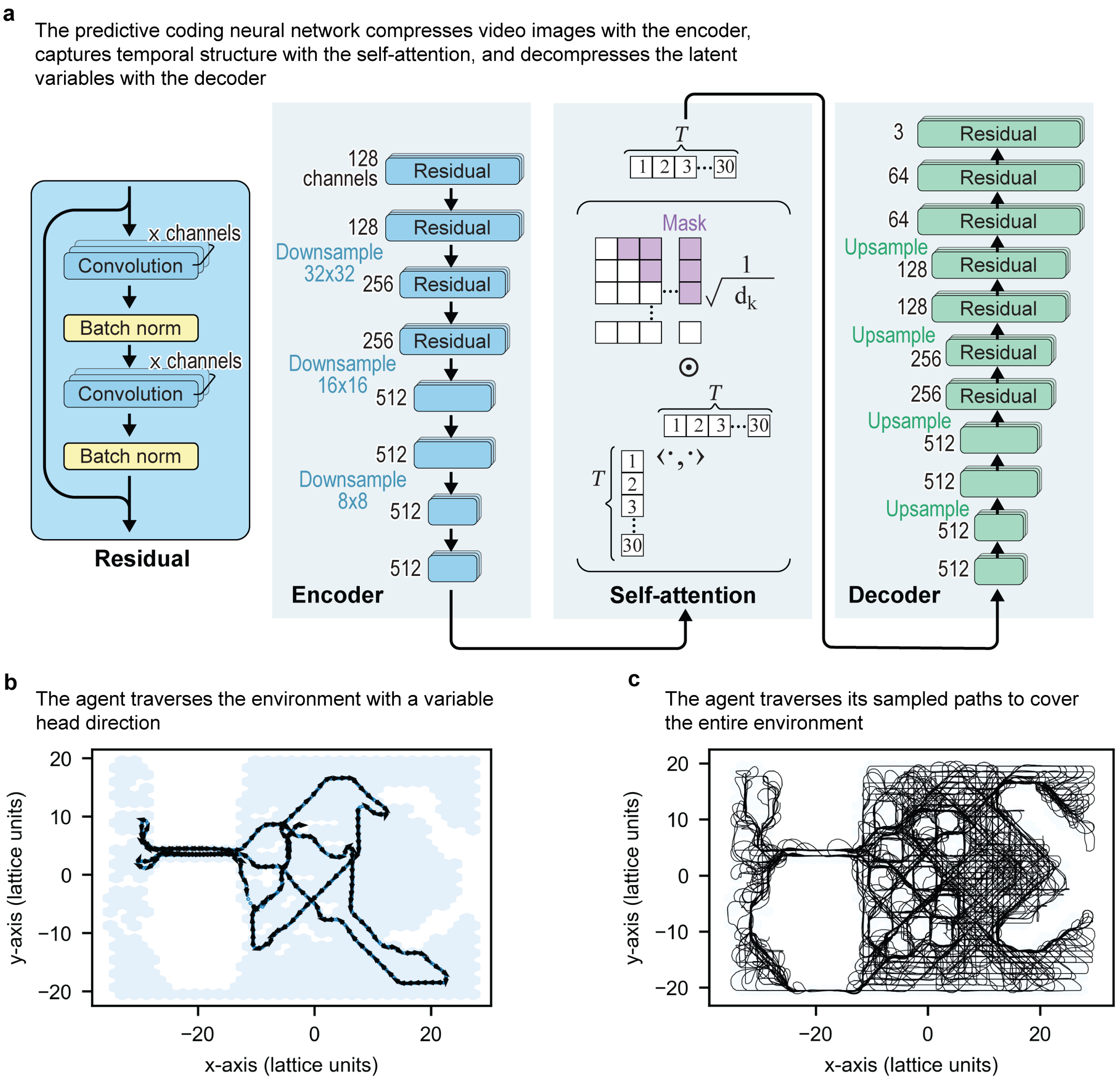


**Figure S5. Extended neural network architecture and training description.** **a,** the predictive coding neural network, or predictive coder, uses an encoder, self-attention, and decoder to perform predictive coding. The encoder is a convolutional neural network architecture called ResNet-18 that uses residuals to compress the high-dimensional video image. The self-attention module capture temporal dependencies from the low-dimensional encoded images. The self-attention's output gives the predictive coder's latent variables. The decoder is a convolutional neural network that upscales—rather than downscales—the predictive coder's latent variables to predicted images. **b,** an example path of the agent shows the agent traversing the environment with a variable head direction. **c,** the agent's paths traverse the space to cover the entire environment.

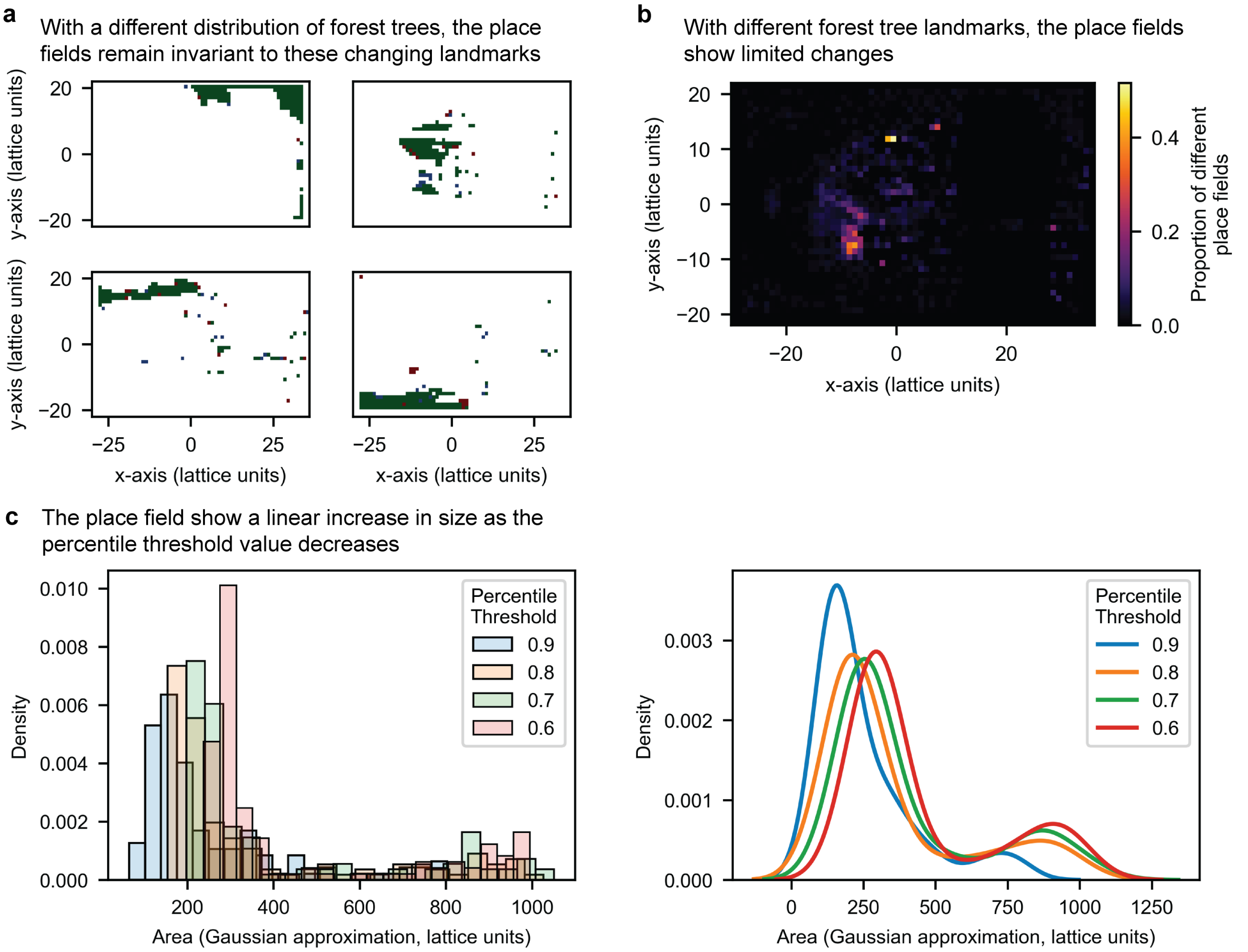


**Figure S6. Extended place field results. a-b,** The predictive coder's place field latent variables are invariant to shifting landmarks. To determine the effect of shifting landmarks, the trees in the environment were removed and randomly redistributed in the forest region. **a,** the predictive coder's original place fields (**red**) were overlaid with the new place fields (**blue**), and the union set of the original and new place fields are shown in green. The new, shifted landmark place fields demonstrate a large overlap (Jaccard index ($|A \cap B|/|A \cup B|$) = 0.828) with the original place fields. **b,** a visual overlap of the proportion of different place fields at every location. The place fields show no variability outside the forest and low variability inside the forest region. **c,** the place fields are measured by thresholding the predictive coder's latent variable, so the place field sizes are dependent on the percentile threshold value. To determine the effect of the threshold value on the place field sizes, the place field size histogram (left) is plotted with respect to the percentile threshold value, and the place field size densities (right) are estimated using kernel density estimation from the histogram.
